# Planarian CREB-binding protein (CBP) gene family regulates stem cell maintenance and differentiation

**DOI:** 10.1101/2020.09.08.287045

**Authors:** Susanna Fraguas, Sheila Cárcel, Coral Vivancos, Ma Dolores Molina, Jordi Ginés, Judith Mazariegos, Thileepan Sekaran, Kerstin Bartscherer, Rafael Romero, Francesc Cebrià

## Abstract

The regulation of stem cells plasticity and differentiation is still an open question in developmental biology. CBP (CREB-binding protein)/p300 is a conserved gene family which functions as a transcriptional co-activator and shows an important role in a wide range of cellular processes, such as cell death, DNA damage response and tumorigenesis. Moreover, CBPs have an acetyl transferase activity that is relevant as histone and non-histone acetylation results in changes in chromatin architecture and protein activity that affects gene expression. Many studies have shown the conserved functions of CBP/p300 on stem cell proliferation and differentiation. The planarian *Schmidtea mediterranea* is an excellent model to study *in vivo* the molecular mechanism underlying stem cell differentiation during regeneration. We have identified five different *Smed-cbp* genes in *S. mediterranea* that show different expression patterns. Functional analyses indicate that *Smed-cbp-2* seems to be essential for stem cell maintenance and cell survival. On the other hand, the silencing of *Smed-cbp-3* results in the growth of apparently normal blastemas; however, these remain largely depigmented and undifferentiated. *Smed-cbp-3* silencing affects the differentiation of several cell lineages including neural, epidermal, digestive and excretory cell types. Finally, we have analyzed the predicted interactomes of CBP-2 and CBP-3 as an initial step to better understand their function on planarian stem cell biology.

## INTRODUCTION

How stem cells are maintained and differentiate into all their distinct lineages is still an open question. Animals capable of regenerating offer us the opportunity to study stem cell behaviour *in vivo* in a context where they are used to rebuild tissues, organs or large and complex body parts. Among those models capable of whole-body regeneration, freshwater planarians have the attractiveness to rely on a population of pluripotent adult stem cells, called neoblasts (Baguñà, 2012; Rink, 2013; Reddien, 2018). After any type of incision or amputation in their body, a muscle contraction takes place in just a few minutes to close the wound (Chandebois, 1980; Baguñá et al., 1988). Later, a quick wound response followed by a later regenerative one will trigger a programme that activates neoblast proliferation and differentiation together with cell death and the re-establishment of the proper polarity (Saló and Baguñà, 1984; Iglesias et al., 2008; Molina et al., 2007, 2011; Gaviño et al., 2011; Pellettieri et al., 2010; Wenemoser and Reddien, 2010; Sandmann et al., 2011; Wenemoser et al., 2012; Owlarn et al., 2017). In recent years, several studies have proved that neoblasts constitute a very complex and heterogeneous cell population (Scimone et al., 2014; Zhu et al., 2016). Thus, a small percentage of neoblasts, the so-called cNeoblasts (clonogenic neoblasts) are real pluripotent stem cells based on the fact that when transplanted individually into irradiated planarians (and therefore completely depleted of neoblasts) are capable to repopulate the host and differentiate into all of the planarian cell types (Wagner et al., 2011). Recently, these cNeoblasts have been further characterized and found to express a tetraspanin gene that has become the first specific molecular marker for the pluripotent neoblasts (Zeng et al., 2018). However, little is known about how neoblasts are directed to differentiate into their multiple lineages (Zhu et al., 2015; Barberán et al., 2016). Post-translational modifications, chromatin remodelling and epigenetic regulation play an important role in stem cell biology (Avgustinova and Benitah, 2016, Godini et al., 2018, Wang et al., 2014). Interestingly, in planarians, some studies have uncovered the role of histone and non-histone post-translational modifications on neoblast biology (Dattani et al., 2019; Strand et al., 2019). Thus, in *Schmidtea mediterranea* the silencing of different components of the nucleosome remodeling and deacetylase (NuRD) complex as *Smed-CHD4* (Scimone et al., 2010), *Smed-HDCA-1* (Robb and Sánchez Alvarado 2014), *Smed-RbAP48* (Bonuccelli et al., 2010) and *Smed-p66* (Vásquez-Doorman and Petersen 2016) results in defects in neoblast proliferation and differentiation. The inhibition of ubiquitination and SUMOylation impairs normal neoblast biology and regeneration (Henderson et al., 2015; Strand et al., 2018; Thiruvalluvan et al., 2017). Also, it has been reported that the planarian homologues of the COMPASS family of MLL3/4 histone methyltransferases have an important role on neoblast proliferation and differentiation and play a conserved role as a tumor suppressor gene in these animals (Mihaylova et al., 2018). Finally, a conserved presence of bivalent promoters in planarian neoblasts has been suggested (Dattani et al., 2018).

The conserved CBP/p300 family is composed of two related transcriptional co-activating proteins: CREB-binding protein (CBP) and p300 (Lundblad et al., 1995), which interact with numerous transcription factors to regulate the expression of their target genes. CBP/p300 can achieve their function by acetylating histones and non-histone proteins as well as serving as scaffold proteins to bring together different factors within the promoter regions. They are members of the lysine acetyltransferase type 3 (KAT3) family (Dutto et al., 2018), which is present in many organisms such as mammals, worms, flies and plants (Yuan and Giordano, 2002). This family interacts with many cellular signalling pathways, such as Notch (Brai et al., 2015), NFkB (Wen et al., 2010), calcium (Hardingham et al., 2001) or TrkB signalling (Esvald et al., 2020), carrying out its function by interacting with numerous transcription factors and other regulatory proteins (Bedford et al., 2010). It has been demonstrated that the CBP/p300 family protein is involved in many cellular processes, including cell self-renewal, proliferation, survival, differentiation, synaptic plasticity, DNA damage response and cell cycle regulation (Shaywitz and Greenberg, 1999; Giordano and Avantaggiati, 1999; Mayr and Montminy, 2001; Goodman and Smolik, 2000; Manegold, P. et al., 2018; Chan and La Thangue, 2001). Here we have identified and characterized five *cbps* homologue genes in *S. mediterranea*. Whereas *Smed-cbp-2* seems required for stem cell maintenance, *Smed-cbp-3* is necessary for the differentiation of several cell lineages, including the central nervous system (CNS), epidermal, gut and protonephridia. Based on the known interactions between CBP/p300 proteins and different factors in humans we have used PlanNET (Castillo-Lara and Abril, 2018) to predict the interactome of planarian CBP proteins and identified several interactions that could help to understand the role of these genes in neoblast maintenance and differentiation.

## RESULTS

### CBP/p300 family in Schmidtea mediterranea

Searching through the current available genome (Grohme et al., 2018), five *cbp* homologues were identified in *Schmidtea mediterranea*, and named *Smed-cbp-1, -2, -3, -4* and -*5* . Phylogenetic analysis showed that planarian CBP homologues group together suggesting that they originated from duplications of the *cbp* gene within the Platyhelminthes lineage (Fig. S1). In order to characterize their expression patterns, whole mount *in situ* hybridizations (WISH) were performed in intact and regenerating animals (Fig. 1). In intact planarians, *cbp-1, -2, -3* and *-4* were expressed rather ubiquitously mainly in the mesenchyme and in the central nervous system (CNS)*; cbp-5* was expressed in the mesenchyme around the pharynx. During regeneration *cbp-2* and *cbp-3* were expressed in the newly formed brain primordia from early stages (Fig. 1A). Recently, planarian cell type atlases based on single-cell sequencing have been reported (Plass et al., 2018; Fincher et al., 2018). These resources allow the analysis *in silico* of the expression dynamics of planarian genes along time in cell types belonging to specific cell lineages. Single-cell analyses of the expression of planarian *cbp* genes largely agree with the patterns obtained after WISH (Fig. 1B). For lineages such as muscle, neuronal, pharynx and protonephridia *cbp-1*, *cbp-2* and *cbp-3* display similar dynamics and show increasing levels of expression during the process of differentiation. On the other hand, the high expression of *cbp-4* and *cbp-5* at early stages appear to decay in parallel to the differentiation of progenitor cell types towards the different cell lineages (Fig. S2).

**Fig. 1.**
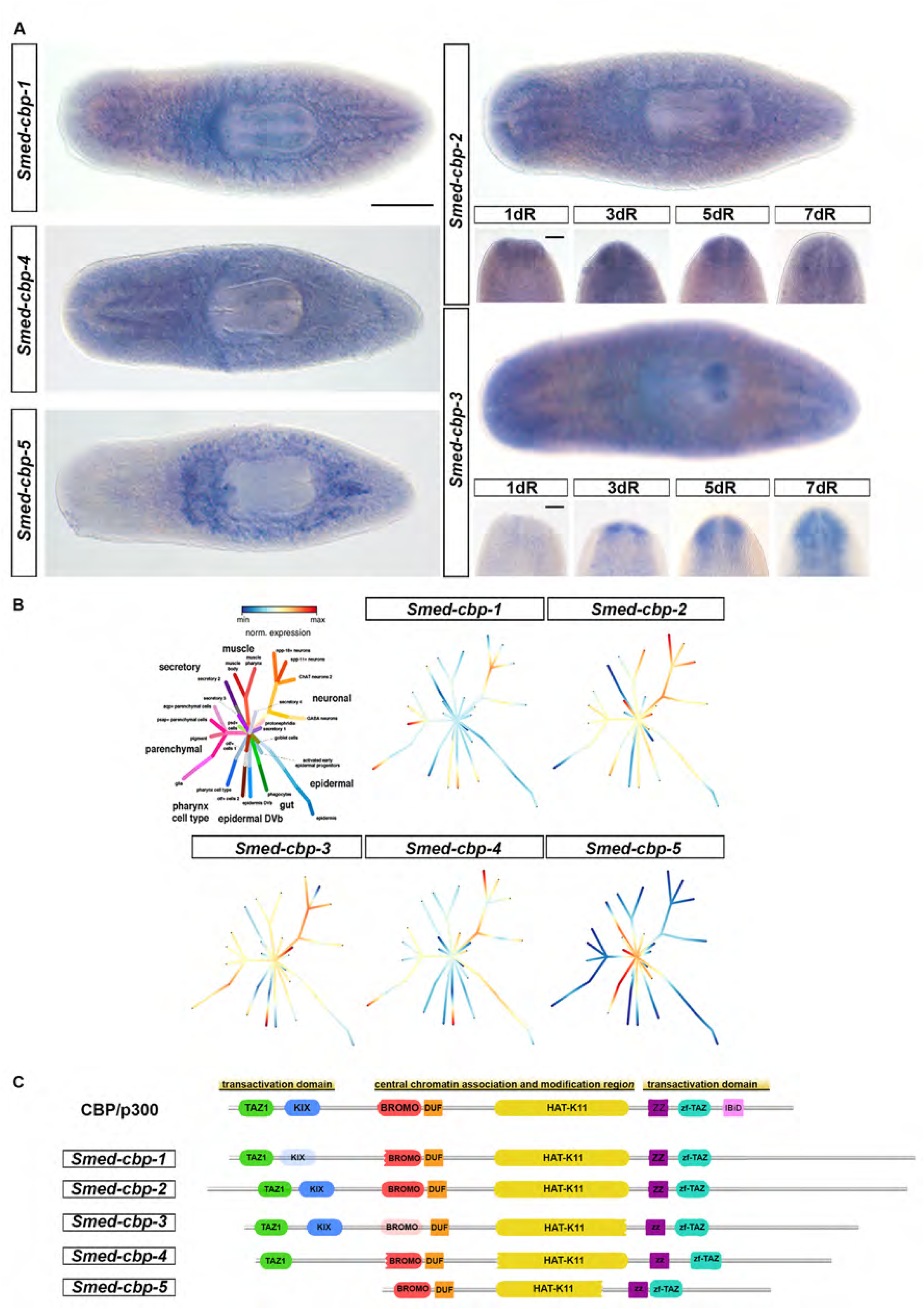
*Schmidtea mediterranea cbp* genes. (A) Expression patterns of *cbp* genes by whole mount in situ hybridization (WISH) in intact (*cbp-1, -4, -5, -2* and -*3*) and regenerating animals (*cbp-2* and -*3*). Scale bar: 400 µm in intact planarians and 200 µm in regenerating animals. Anterior to the left in intact animals and to the top in regenerating animals. (B) Single-cell transcriptomic expression of *Smed-cbp-1, -2, -3, -4* and -*5* along all the planarian cell types. (C) Schematic diagram of the domain arrangement of the different S. mediterranea CBP proteins compared to vertebrate CBP/p300. The different domains were marked in different colours. Shallowed colour of the bromodomain of Smed-CBP-3 and the KIK domain on Smed-CBP-1 denote weak conservation.

CBP and p300 have similar structures and share a modular organisation with five protein interaction domains (Fig. 1C; Wang et al., 2013). In the middle of the sequence there is a central chromatin association and modification region, which encompasses the lysine acetyltransferase domain HAT-K11 and the bromodomain. Thanks to its lysine acetyltransferase activity, CBP and p300 are able to acetylate both histones and non-histone factors (Dancy and Cole, 2015). The central region is flanked by a characteristic structure, composed of several transactivation domains: a transcriptional-adaptor zinc-finger domain 1 (TAZ1); a kinase inducible domain of CREB interacting domain (KIX); a domain of unknown function (DUF); a zz-type zinc finger domain (ZZ); a transcriptional-adaptor zinc finger domain 2 (zf-TAZ) and a IRF3-binding domain (IBiD). *S. mediterranea* CBP proteins contain most of these essential conserved domains (Fig. 1C).

### *cbp-2* is required for regeneration and tissue homeostasis

To characterize the function of *Smed-cbp* genes we performed RNA interference (RNAi)-based functional analyses . No obvious defects were observed after silencing *cbp-1*, *-4* and *-5* (Fig. S3). On the contrary, the silencing of *cbp-2* and *cbp-3* significantly impaired the viability or the regeneration of the treated animals.

*cbp-2* RNAi treated animals resulted in a failure to form a proper blastema (Fig. 2A) and all animals died in less than 15 days after amputation. After silencing *cbp-2,* the treated planarians did not regenerate a normal blastema at 3 of regeneration compared to controls. On day 5, control animals had already differentiated the eyes, whereas *cbp-2*(RNAi) animals did not show any sign of eye differentiation. After 7 and 10 days of regeneration, *cbp-2*(RNAi) animals differentiated smaller and aberrant eye-pigment cups within very small blastemas and closer to the pre-existing post blastema region (Fig. 2A). The efficiency of the RNAi silencing was measured by qPCR after 7 days of regeneration (Fig. 2B).

**Fig. 2.**
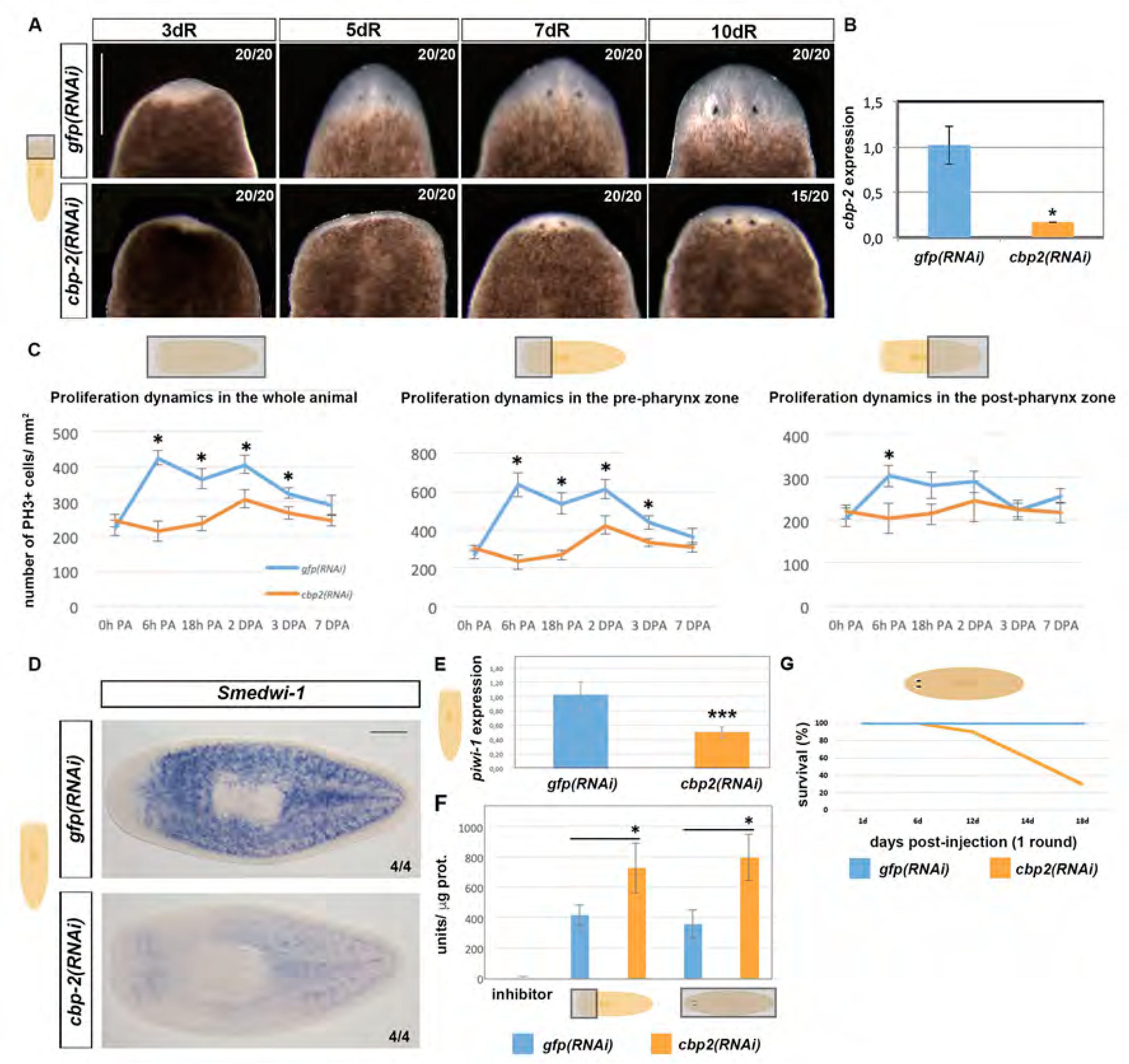
*cbp-2* is required for regeneration and survival. (A) Alive *cbp-2* and *gfp*(RNAi) animals. *cbp-2*(RNAi) regenerating trunk pieces fail to form a proper blastema but do differentiate eyes. Scale bar: 300 µm (B) *cbp-2* qPCR expression analysis after *gfp* and *cbp-2*(RNAi) at 7days of regeneration. *cbp-2* expression is strongly downregulated after *cbp-2* inhibition (*P< 0,05). (C) Quantification of mitotic PH3^+^ immunolabeled cells in *gfp*(RNAi) and *cbp-2*(RNAi) animals at several time points post amputation (DPA) reveals a reduced number of mitoses after silencing *cbp-2* (*P< 0,05, Student’s t-test. Values represent the mean ± s.e.m of an average of 7 samples per time point and condition). (D) *Smedwi-1* WISH at 7 days or regeneration (7dR). *cbp-2* silencing strongly decreases the neoblast population. Scale bar: 400 µm. (E) *Smedwi-1* expression quantified by qPCR analysis is significantly decreased after *cbp-2*(RNAi) at 7dR, (F) Caspase-3 activity quantification in *cbp-2(RNAi)* and control animals, in both regenerating and intact treated planarians at 8 days post amputation or post-injection, respectively (*P< 0,05). (G) Percentage of survival after *cbp-2* silencing.

In order to check if cell proliferation was affected after silencing *cbp-2,* we carried out an immunostaining with the anti-PH3 antibody in control and *cbp-2*(RNAi) animals. *S. mediterranea* presents two mitotic peaks during the regenerative response: the first mitotic peak is global and takes place at 6 hours after amputation; the second mitotic peak is seen at 48 hours after amputation and is restricted to a narrow region of the post-blastema adjacent to the wound region (Saló and Baguñà, 1984; Wenemoser and Reddien, 2010). We quantified the number of mitoses along several time points after head amputation (Fig. 2C). Although mitotic neoblasts were detected after *cbp-2* RNAi a significant reduction of proliferating cells was detected in all time points.

The first mitotic peak was not detected after silencing *cbp-2*; however, the second peak at 48h appears to occur although significantly attenuated compared to controls. WISH with the neoblast marker *Smedwi-1* revealed a strong decrease in the neoblast population at 7 dpa, compared to controls, suggesting an important role of *cbp-2* for the maintenance of the stem cell population (Figure 2D). A decrease in the expression of *Smedwi-1* was also observed by qPCR (Fig. 2E). Concomitant to the decrease in the number of proliferating cells, *cbp-2*(RNAi) animals showed an increase in cell death in a caspase-3 activity assay (Fig. 2F). The silencing of *cbp-2* in intact non-regenerating animals also led to their death in 2-3 weeks (Fig. 2G). In intact animals an increase of cell death was also observed (Fig. 2E). Overall, these results indicate that *cbp-2* is required for survival and suggest that could have an important role in the maintenance of the neoblast population and cell survival.

### Differentiation and patterning defects after *cbp-2* RNAi

As it was observed that even in the presence of extremely reduced blastemas eye-pigment cups were differentiated (Fig. 2A) we analysed in more detail the differentiation of several cell types and tissues in the *cbp-2(RNAi)* animals. Regenerating head pieces seemed to normally extend new posterior gut branches and ventral nerve cords, as well as to form a pharynx cavity (Fig. 3A). On the other hand, most of the *cbp-2* RNAi animals failed to differentiate new pharynges (Fig. 3A). During anterior regeneration, the differentiation of the digestive and excretory system within the blastema was not apparently affected (Fig. 3B). In contrast, the regeneration of the normal pattern of visual projections was impaired. Although *cbp-2(RNAi)* animals were able to differentiate new photoreceptor cells, they could not extend visual axons to form a proper optic chiasm as controls (arrowhead in Fig. 3B). Interestingly, brain tissue differentiated in the wound region of *cbp-2(RNAi)* animals (Fig. 3C). However, and similar to what was observed for the photoreceptors, the new brain cells did not form properly patterned cephalic ganglia; the two small brain rudiments were not connected by a transverse commissure and appeared mainly in the pre-existing post-blastema region. Moreover, whereas in controls mitotic cells were normally found behind the new cephalic ganglia, after silencing *cbp-2* proliferating neoblasts were detected at the tip of the head in front of the newly differentiated brain tissues (arrowhead in Fig. 3C). Even though patterning defects were observed for several tissues, *cbp-2(RNAi)* animals showed normal expression of posterior and anterior polarity markers (Fig. S4).

**Fig. 3.**
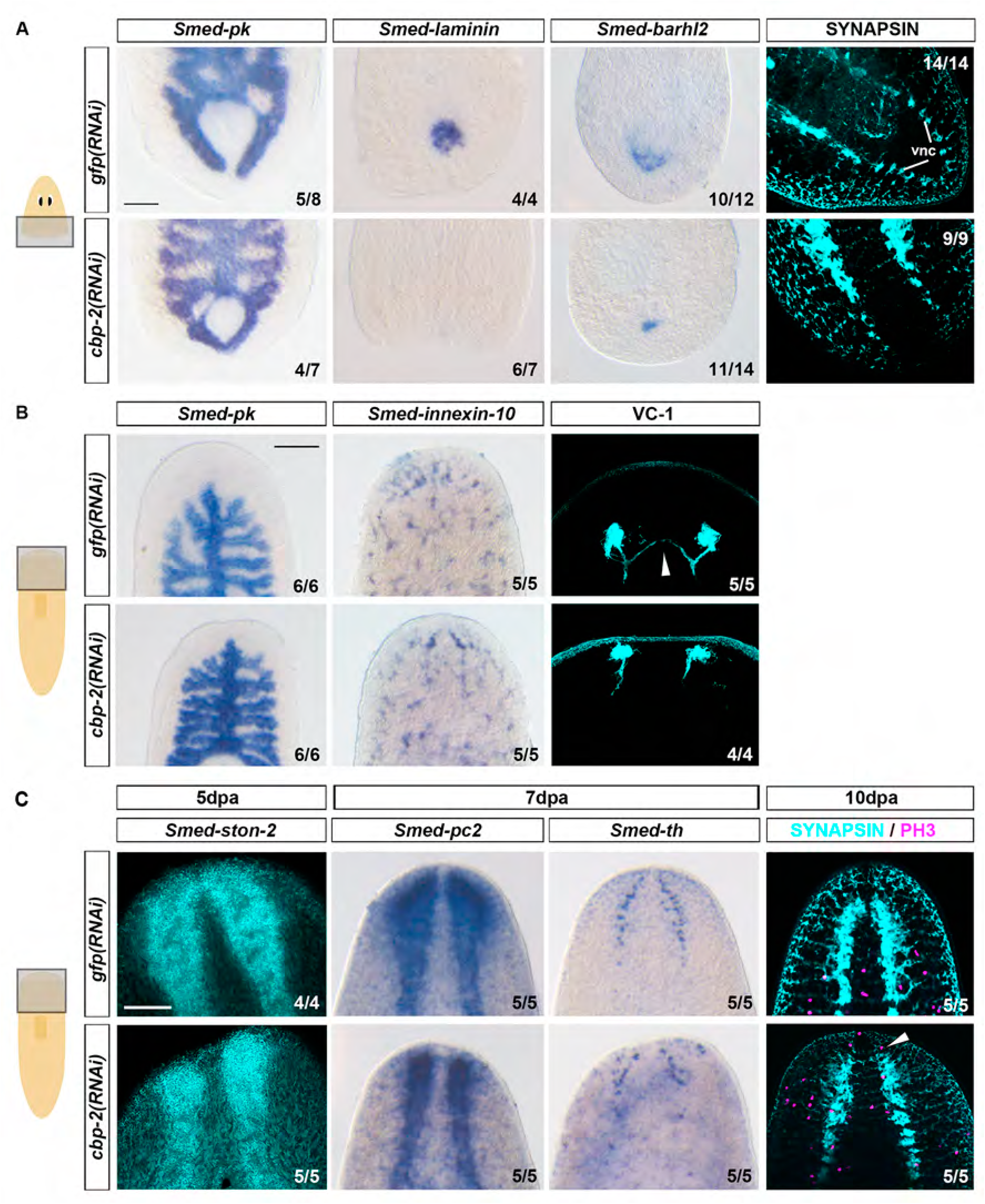
Tissue and patterning regeneration defects after silencing *cbp-2*. (A) Posterior gut regeneration impaired after silencing *cbp-2*. Pharynx formation is also inhibited as seen with the markers *Smed-laminin*, *Smed-barhl2* and anti-SYNAPSIN. Posterior ventral nerve cords (vnc) regeneration is impaired after silencing *cbp-2*, as seen with the anti-SYNAPSIN. Samples correspond to regenerating heads after 7 days of regeneration. Scale bar: 200 µm. (B) Digestive and excretory system regeneration, visualized with *Smed-pk* and *Smed-innexin-10*, respectively, is not affected after silencing *cbp-2* in regenerating trunks after 7 days of regeneration. However, the visual system visualized with VC-1 shows an abnormal pattern with no optic chiasm (arrowhead) formed. Scale bar: 200 µm. (C) CNS regeneration, visualized with *Smed-ston-2, Smed-pc2, Smed-th* and anti-SYNAPSIN, was strongly disrupted in *cbp-2* RNAi regenerating trunks after 5-10 days of regeneration. Moreover, these treated animals show an abnormal anterior distribution of mitoses (white arrowhead). Scale bar: 200 µm. In all images, anterior to the top.

### *cbp-2* is required for the maintenance of the epidermal lineage

We next performed WISH for several epidermal markers to determine the role of *cbp-2* during epidermal maturation. Planarian epidermis is a monostratified tissue composed by a single layer of both non-ciliated and multi-ciliated differentiated cell types (Rompolas et al., 2010). The epidermal lineage is well characterized in planarians (Zhu et al., 2015; Cheng et al., 2018; Zhu and Pearson, 2018). Epidermal maturation requires temporally correlated transition states in which lineage-commitment zeta neoblasts (*Smed-zfp-1*+, van Wolfswinkel et al., 2014) become post mitotic and start to sequentially express *Smed-NB.21.11e* (“early progeny progenitor cells”) and *Smed-AGAT-1* (“late progeny progenitor cells”) and finally differentiate in mature epidermal cells that express genes such as *ifb, rootletin* and *PRSS12* (Wurtzel et al., 2017). Knockdown of *cbp-2* RNAi resulted in an evident reduction of *zfp-1+* cells (Fig. 4). Concurring with these results, we also observed a clear depletion of epidermal progenitor cells after silencing *cbp-2* (Fig. 4). To determine the effects of this decrease in epidermal progenitors on epidermis differentiation, we analyzed the expression of specific markers for the mature epidermis. In control animals, *Smed-PRSS12* is expressed throughout the dorsal and ventral epidermis in both ciliated and non-ciliated epidermal cells, whereas *Smed-rootletin* is only expressed in ciliated epidermal cells (Wurtzel et al., 2017). After silencing *cbp-2*, the expression of both genes was remarkably decreased. Similarly, anti-TUBULIN immunostaining revealed less and disorganized pattern of cilia, compared to control animals. However, on the other hand, *ifb*+ mature epidermal cells appeared to differentiate normally along the dorsal-ventral boundary of the newly formed blastema in *cbp-2* RNAi animals (Fig. 4).

**Fig. 4.**
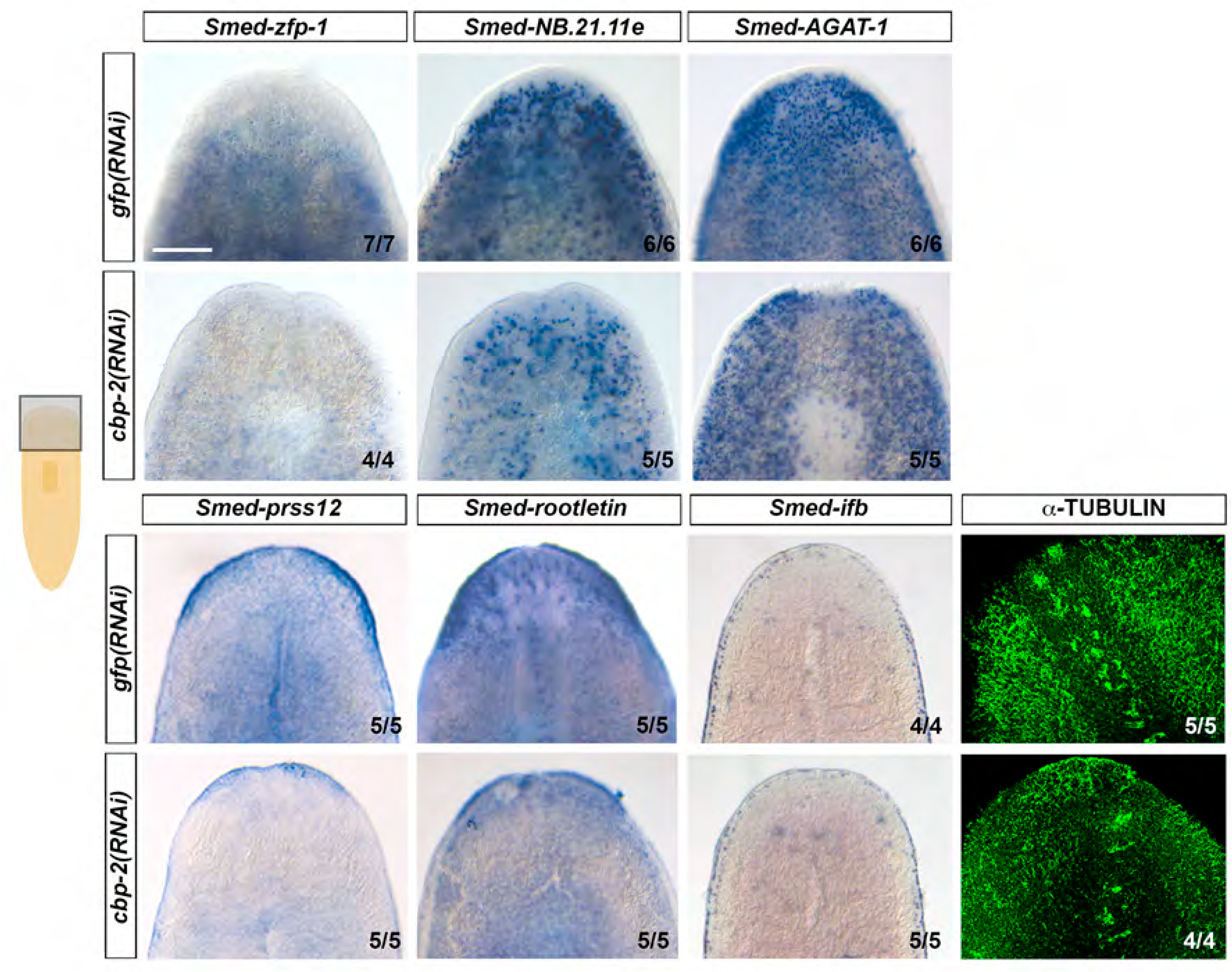
Defects in the epidermal lineage after silencing *cbp-2*. *cbp-2* RNAi causes an evident decrease of ζneoblasts (*zfp-1+)* and early and late stage epidermal progenitors (*Smed-NB.21.11e* and *Smed-AGAT-1*, respectively). No differences in the expression of *ifb+* in the epidermal cells along the dorsoventral margin showed no differences between control and *cbp-2* RNAi animals are seen, less expression of *Smed-rootletin* and *Smed-prss12* in the differentiated epidermal cells is clearly observed after *cbp-2* silencing. Anti-TUBULIN immunostaining shows less and disorganized cilia in *cbp-2* RNAi animals. All samples correspond to 7 days regenerating trunks. Scale bar: 200 µm. In all images, anterior to the top.

### Gut and eye progenitor cells decrease after *cbp-2* RNAi

In order to characterize how the decrease in the neoblast population affected the specification of distinct populations of progenitors, double labelling with cell progenitor specific markers and the neoblast marker anti-SMEDWI-1 were carried out (Fig. 5). By using *hnf-4* (Wagner et al., 2011) and *ovo* (Lapan ad Reddien, 2012) as specific markers for gut and eye progenitors, respectively, a significant reduction in the number of these progenitors was observed after silencing *cbp-2* (Fig. 5A, B). On the other hand, the number of *sim+* neural progenitors (Cowles et al., 2013) increased significantly after silencing *cbp-2*, whereas *ston-2+* neural progenitors (Molinaro and Pearson, 2016) showed no significant differences although it also showed a tendency to increase (Fig. 5C). These results suggest that, in agreement with a reduction in the neoblast pool, different populations of cell specific progenitors decrease after the silencing of *cbp-2*. Altogether with the results of the previous section, our data suggest that although reduced (gut and eye) or increased (neural) numbers of progenitors were specified after *cbp-2* RNAi, the new mature cells that differentiate from them cannot give rise to properly patterned tissues and organs.

**Fig. 5.**
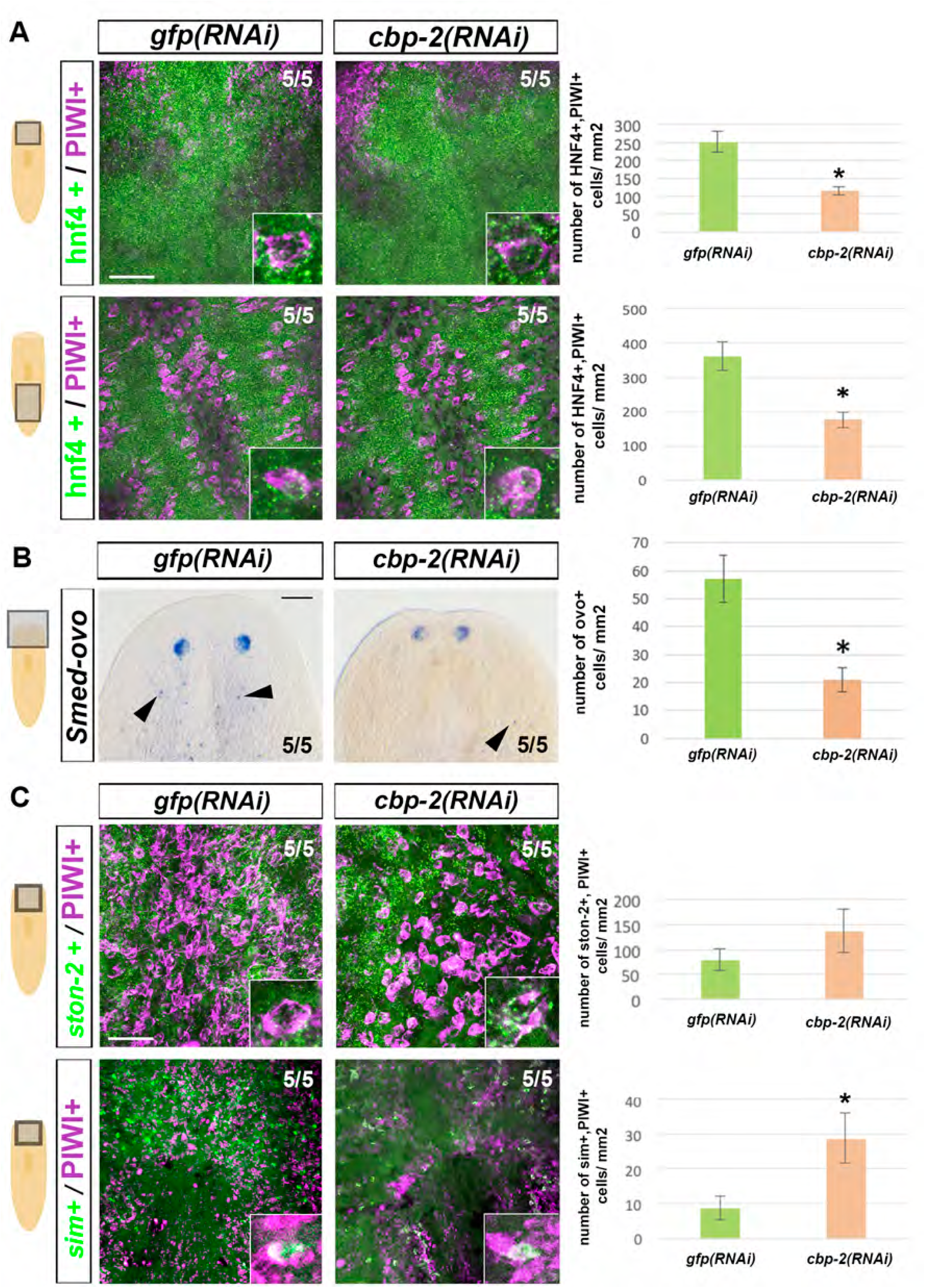
Defects in cell progenitor compartments after *cbp-2* silencing. (A) Double staining and quantification of *Smed-hnf4/* SMEDWI-1 gut progenitors. A magnified view of the progenitors cells is boxed in each panel. Scale bar: 100 µm. (B) Knockdown of *cbp-2* causes a significant decrease of eye progenitors (arrowheads), labelled with *Smed-ovo*. (C) *Smed-sim/SMEDWI-1* neural progenitors labelled significantly increase after silencing *cbp-2*. No significant differences are observed for *Smed-ston-2/SMEDWI-1* progenitors. Scale bar: 200 µm. In all quantifications, *P< 0,05, Student’s test. Values represent the mean ± s.e.m of an average of 5 samples per each condition.

### *cbp-3* is required for blastema differentiation

The silencing of *cbp-3* by RNAi (Fig. 6) resulted in blastemas that grew in size but did not show any external sign of differentiation in terms of eyes or body pigmentation (Fig. 6A). Up to 5 days post amputation, the blastema of the *cbp-3(RNAi)* animals grew at the same rate as in controls. After that, those blastemas kept growing but at a lower rate compared to controls (Fig. 6B). The efficiency of *cbp-3* RNAi was quantified by qPCR (Fig. 6C). Next, the mitotic response of neoblast to amputation was analysed. Remarkably, the first mitotic peak that occurs at 6h as a normal wound response was significantly increased after silencing *cbp-3* (Fig. 6D). Moreover, whereas in controls the first mitotic peak is normally followed by a decrease in the proliferative activity up to 18h when proliferation increases again to reach a second mitotic peak at 48h, in *cbp-3(RNAi)* animals proliferation kept increasing from the amputation time up to 48h. After that, proliferation rate decreased at the same rate as in controls and no further differences were detected (Fig. 6D-E). Previous studies have demonstrated that the blastema is formed by the entry of neoblasts that once there leave the cell cycle and differentiate into the multiple missing cell types (Reddien, 2013; Scimone et al., 2014). As *cbp-3(RNAi)* animals showed blastemas with no signs of external differentiation and neoblast proliferation did not appear to be impaired, the cellular composition of those blastemas was analysed with different neoblast specific markers (Fig. 6F). After 10 days of regeneration, control animals showed a normal distribution of neoblasts, mostly absent within the blastema. In contrast, the blastemas of *cbp-3(RNAi)* animals were full of neoblasts (Fig. 6F) suggesting that the silencing of *cbp-3* impairs neoblast differentiation within them. Finally, the silencing of *cbp-3* in intact animals did not result in any apparent phenotype in the morphology of the gut and the CNS nor in the neoblast proliferative rate (Fig. S5).

**Fig. 6.**
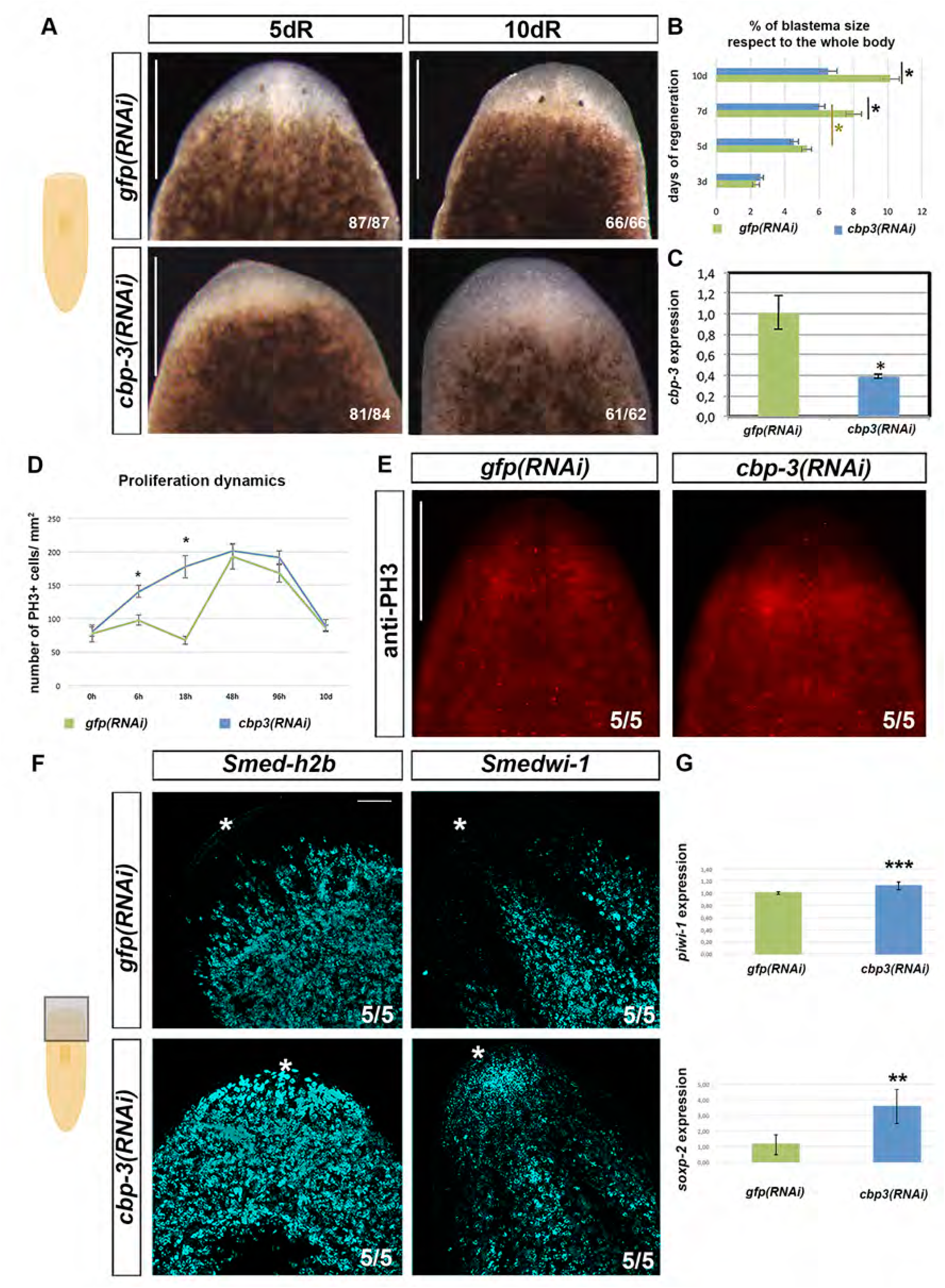
Undifferentiated blastemas after *cbp-3* silencing. (A) *cbp-3* RNAi animals grow unpigmented blastemas with no sign of eye differentiation. Scale bar: 300 µm. (B) Quantification of blastema size measured as the square root of the total non pigmented area, divided by the square root of the total worm area. Black asterisks: comparison between *cbp-3* RNAi and control animals at the same time point; yellow asterisk: comparison between *cbp-3* RNAi animals at different time points (values represent the mean ± s.e.m of an average of 30 samples per time point, Student’s t-test, *P>0,05). (C) Quantification of the efficiency of *cbp-3* RNAi by qPCR (*P< 0,05). (D) Quantification of mitotic cells at several time points of regeneration. (E) Distribution of PH3+ mitotic cells after 10 days of regeneration.. (F) Neoblasts labeled with *Smed-h2b* and *Smedwi-1* accumulate within the blastema of *cbp-3* RNAi animals at 10 days of regeneration. Asterisk marks the tip of the head. qPCR analyses show an increased expression of the neoblast markers *Smedwi-1 and Smed*-*soxp-2* after *cbp-3* RNAi at 7dR. (*P< 0,05). Scale bar: 100 µm. In all images, anterior to the top.

### *cbp-3* RNAi impairs the differentiation of several cell lineages

As *cbp-3(RNAi)* blastemas were full of neoblasts and did not show any external sign of differentiation, specific markers were used to analyse the differentiation of several cell types and tissues within them. Labellings with pan-neural markers such as *Smed-pc2* or SYNAPSIN revealed that the regeneration of new cephalic ganglia was highly impaired after silencing *cbp-3* (Fig. 7A). In agreement with these results, *cbp-3*(RNAi) animals showed a complete absence or a significant reduction of GABAergic (*Smed-gad*), dopaminergic (*Smed-th*) and brain lateral branches (*Smed-gpas*) neurons (Fig. 7A and S6). Moreover, when positive cells for these markers were detected within the blastema they did not show a proper pattern compared to controls (Fig. 7A). In addition to these defects in the neural lineages, *cbp-3(RNAi)* animals also failed to differentiate new excretory (*Smed-innexin-10*) and gut (*Smed-pk*) cells (Fig. 7B) as well as new pharynges (Fig. S6).

**Fig. 7.**
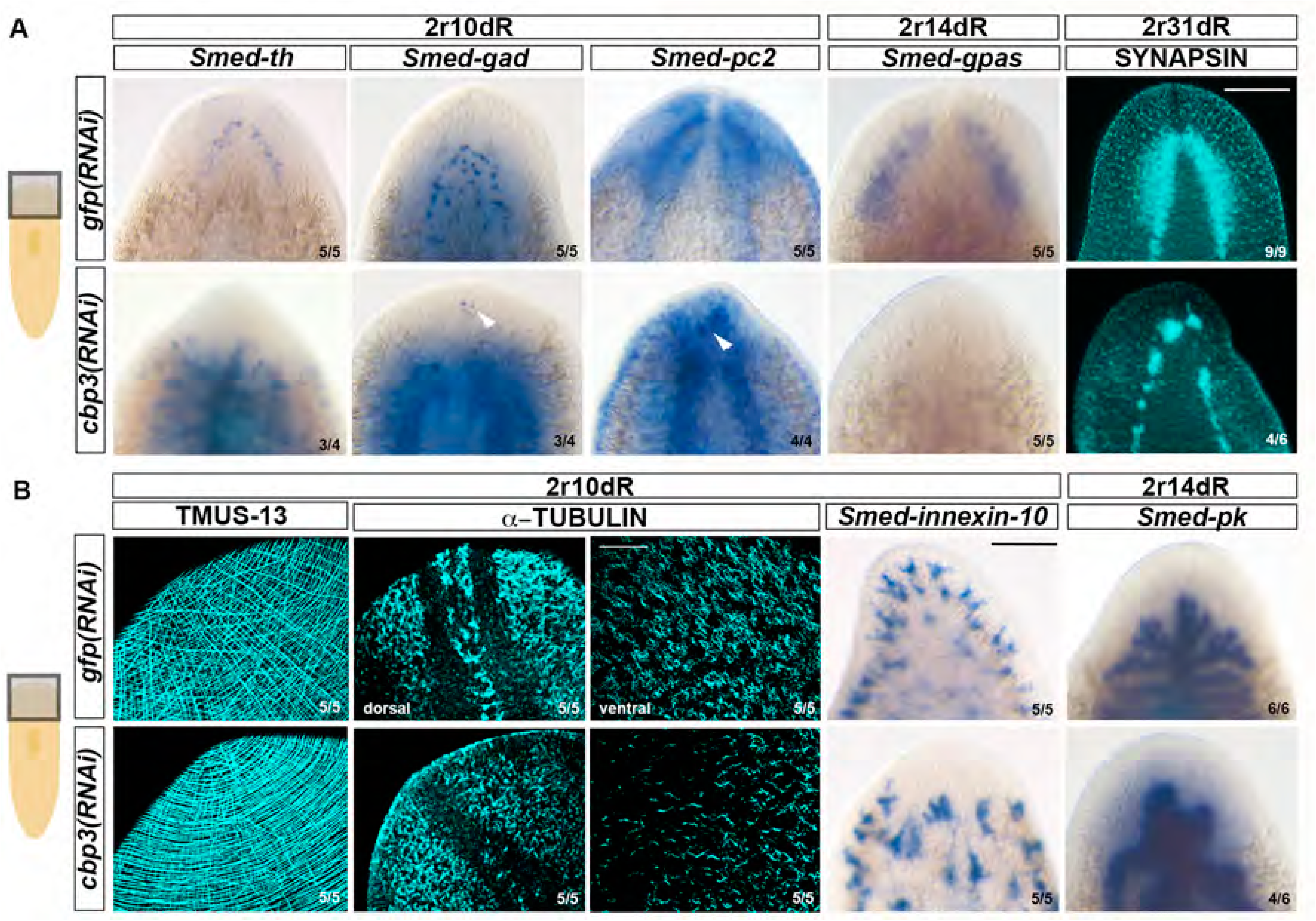
Differentiation defects after silencing *cbp-3* (A) *cbp-3* silencing inhibits CNS differentiation as visualized with several specific markers, checked between 10 and 31 days of regeneration. Scale bar: 200 µm. In all images, anterior to the top. (B) RNA interference of *cbp-3* activity results in normal regeneration of the body-wall musculature labelled with TMUS-13 antibody. Cilia defects in the epidermis, visualized with an anti-TUBULIN antibody in 10 days regenerating fragments. Protonephridial cells labeled with *Smed-innexin-10* are absent of the *cbp-3(RNAi)* blastemas. The anterior gut branch (*Smed-pk*) also fails to differentiate after *cbp-3* silencing. Scale bar: 100 µm for TMUS-13, TUBULIN and SYNAPSIN, and 200 µm for the other images.

On the other hand, the silencing of *cbp-3* did not apparently impaired the normal regeneration of the body-wall musculature as circular, longitudinal and diagonal muscle fibers differentiated in those blastemas (Fig. 7B). Also, epidermal cells differentiated within the blastemas of *cbp-3(RNAi)* animals. Planarian epidermal cells are multiciliated cells (Hyman, 1951) and show distinct patterns in dorsal and ventral surfaces. Dorsally, ciliated epidermal cells are mainly distributed along the midline and in the lateral sides of control animals; ventrally, these epidermal ciliated cells show a uniform distribution (Fig. 7B). In contrast, *cbp-3(RNAi)* animals appeared to regenerate epidermal cells with a reduced number of cilia and lacked the typical dorsal stripe (Fig. 7B). Overall, these results indicate that the differentiation of neural, gut and excretory cells was severely affected after *cbp-3* RNAi; the body-wall musculature, however, was properly regenerated and new epidermal cells appeared within the blastema but showed defects in their cilia.

### Impaired specification and differentiation of cell progenitors after *cbp-3* RNAi

In order to determine whether the defects in cell differentiation were caused by a lack or reduction in the number of lineage-committed progenitors or by a problem in their differentiation to mature cells, the expression of different progenitor specific markers was analysed (Fig. 8). Concerning the epidermal lineage, *Smed-zfp1* was expressed through the mesenchyme except in the most anterior part of the head in control animals, whereas *cbp-3(RNAi)* animals showed a large number of *Smed-zfp-1^+^* cells up to the tip of the head (Figure 8A), in agreement with what it has been shown as the blastemas of these animals are full of neoblasts. Concomitant with this increase in *Smed-zfp-1+* neoblasts, a significant increase in the number of *Smed-NB.21.11e, AGAT-1* and *vimentin* positive cells was also observed (Fig. 8A). Also, whereas a higher expression of *Smed-PRSS1/2* was seen, *Smed-rootletin* seemed to be expressed at lower levels (Fig. 8A) Remarkably, when the epidermal cell density was quantified a significant increase in the number of cells in the epìdermal monolayer was seen compared to controls (Fig. 8A). However, and based on the absence of *ifb+* cells along the dorsoventral border (Fig. 8A) and the defects observed in the cilia (Fig. 7B) these results suggest that *cbp-3* could be required for the final maturation of the epidermal cells (Fig. 8B). A similar analysis was carried out for *Smed-hnf4+* gut progenitors. Also in that case, an increase in gut progenitors was observed after silencing *cbp-3* (Fig. 8C), although no proper gut regeneration was observed (Fig. 7B and S6), suggesting that, similarly to the epidermal lineage, *cbp-3* would be required for the differentiation of the gut progenitors but not for their specification.

**Fig. 8.**
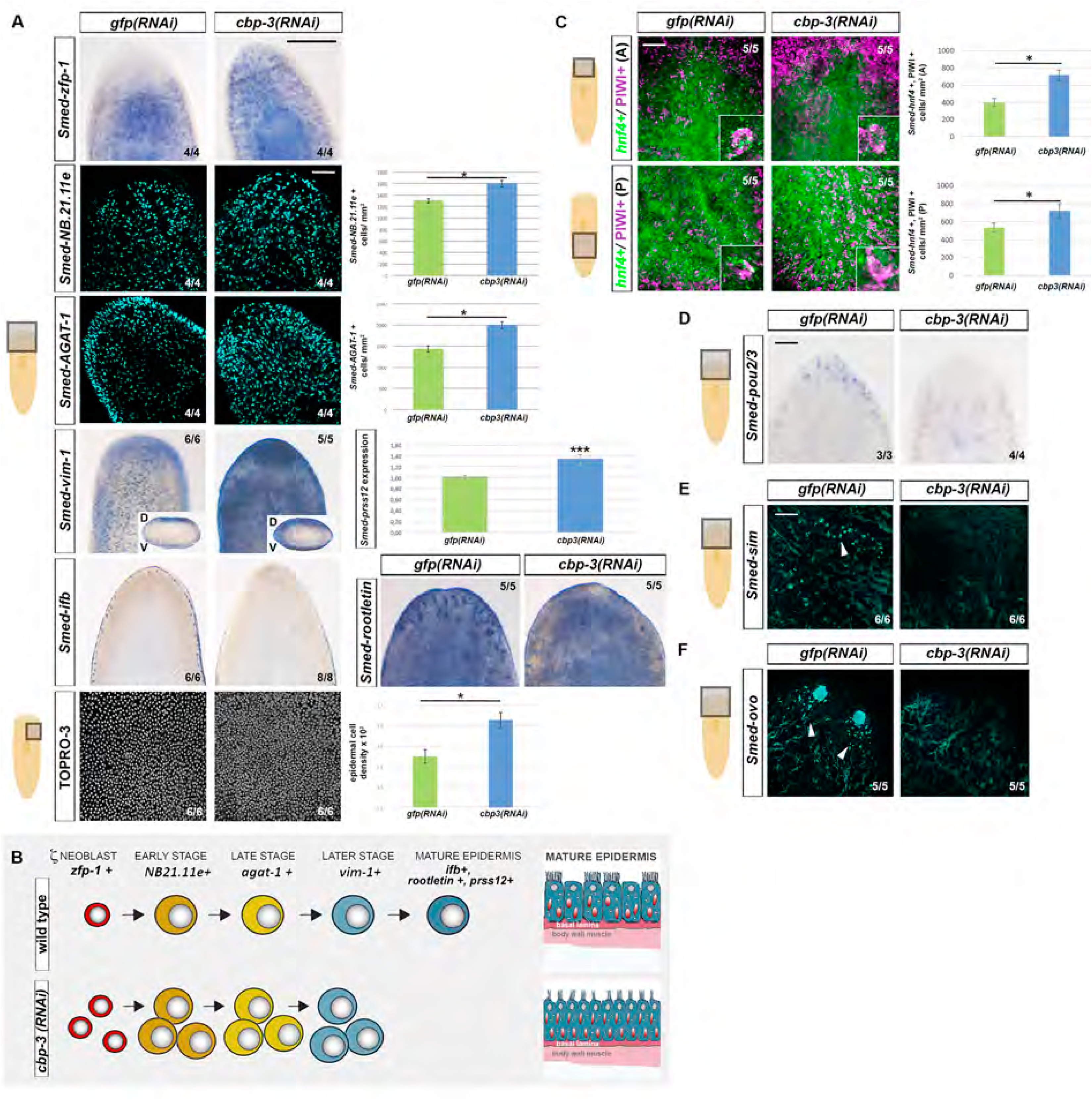
Defects in the cell progenitor compartments after silencing *cbp-3*. (A) The number of epidermal progenitors, labelled with *Smed-zfp-1, NB.21.11.e* and *AGAT-1*, increases significantly after *cbp-3* knockdown. *Smed-vim-1*+ cells and *Smed-prss12* expression quantified by qPCR significantly increase after silencing *cbp-3*. *Smed-ifb* and Smed-rootletin expression decreases after *cbp-3* knockdown. Scale bar: 300 µm for WISH images and 100 µm for WFISH images. Epidermal nuclear density increases after silencing *cbp-3*, as visualized with TO-PRO®-3. Scale bar: 100 µm (B) Scheme summarizing the main defects along the epidermal lineage after *cbp-3* silencing. (C) *Smed-hnf-4*/SMEDWI-1 gut progenitors significantly increase in *cbp-3(RNAi)* animals. (D) *Smed-POU2/3*/SMEDWI-1 protonephridial progenitors are absent for the blastemas of *cbp-3*(RNAi) animals. (E) *Smed-sim* neural progenitors (arrowheads) are practically absent after silencing *cbp-3*. (F) *Smed-ovo* eye progenitors (arrowheads) are practically absent in *cbp-3* RNAi animals. Scale bar in C, D, E, and F: 100 µm. All images correspond to samples at 10 days of regeneration.

On the other hand, the analyses of the expression of excretory and neural progenitors revealed the complete absence of them after silencing *cbp-3* (Fig. 8D-F). Thus, there is a total absence of cells expressing the protonephridial progenitor marker *Smed-POU2/3* (Scimone et al., 2011) after silencing cbp-3 compared to controls (Fig. 8D). Moreover, whereas controls displayed *Smed-sim+* neurons and neural progenitors, as well as *Smed-ovo+* mature photoreceptors and eye progenitors, these markers were totally absent within the blastemas of the *cbp-3(RNAi)* animals (Fig. 8E, F). Also, *Smed-sim+* neurons were practically absent in the mesenchyme of intact planarians after silencing *cbp-3* (Fig. S5). These results agree with the strong impairment in CNS regeneration (Fig. 7A) and suggest that *cbp-3* is required for neural progenitor specification. Taken together these results indicate that, during regeneration, *cbp-3* would have different roles for progenitor specification and differentiation depending on the specific lineage.

### Smed-CBP-2 and Smed-CBP-3 co-expression and interaction with other factors

The results described on the previous sections suggest that after duplication planarian CBPs diverged functionally. Smed-CBP-2 appears mainly required for correct neoblast maintenance and proliferation, whereas Smed-CBP-3 regulates commitment of the stem cells into specific progenitor lineages as well as their final differentiation into well-patterned differentiated tissues. In order to deep insights into these diverse functions, and taking advantage of the available planarian single cell data (Plass et al., 2018) and the recently developed Gene Co-expression Counts tool (Castillo-Lara and Abril 2018; Castillo-Lara et al., 2020), we analyzed in deep the expression of *Smed-cbp-2* and *Smed-cbp-3* in the main planarian cell lineages (Fig. S7).

Our analysis revealed that the relative percentage of *Smed-cbps* expressing cells was considerably uniform among lineages, being *Smed-cbp-3* always more abundant than *Smed-cbp-2*. Precisely, *Smed-cbp-2* expression in the different lineages ranges from 4,4% of the muscle cells to 6,3% of the neuronal cells (Fig. S7A). On the other hand, *Smed-cbp3* expression at the single-cell level ranges from 7% of the epidermal cells of the dorsoventral boundary to 15% of the excretory cells. Both progenitor (*piwi1+*) and differentiated (*piwi1-*) cells of the neuronal, epidermal, muscular and intestinal compartments expressed *Smed-cbps* (Fig. S7B). Interestingly, in most cellular lineages, *Smed-cbps* expression seemed slightly enriched in the progenitor compartment. In particular, in the neural lineage nearly 11% and 13 % of neural progenitor cells but only 5,2 % and 9,1% of the differentiated neural cells expressed *Smed-cbp-2* and *Smed-cbp-3*, respectively. As already mentioned, planarian cell type atlases based on single-cell sequencing display similar dynamics of expression for *Smed-cbp-2* and *Smed-cbp-3* during the differentiation of most planarian cellular types (Fig. 1A-B, S2 and S7B). Remarkably, however, we observed that only a reduced percentage of cells (less than 1% of the cells in most cell lineages) co-express planarian *Smed-cbp-2* and *Smed-cbp-3* genes, representing 9% (175 cbp-2^+^cbp-3^+^ versus 1941 cbp-2^+^cells) and 15% (175 cbp-2^+^cbp-3^+^ versus 1162 cbp-2^+^cells) of the cells (Fig. S7A). Altogether these analyses revealed that *Smed-cbp-2* and *Smed-cbp-3* are barely co-expressed.

CBPs are multifunctional transcriptional co-activators and their diverse functions rely in part on their extended network of protein interactors (Bedford et al., 2010; https://thebiogrid.org; https://string-db.org). To investigate whether the diverse functions of planarian Smed-CBP-2 and Smed-CBP-3 may relate to a diversified network of protein interactors, we took advantage of the tool recently developed by Castillo-Lara and Abril (2018) that predicts planarian protein-protein interactions using sequence homology data and a reference Human interactome (Castillo-Lara and Abril, 2018) and examined the interactome of planarian Smed-CBP-2 and Smed-CBP-3. Surprisingly, regardless of their diversified function, similar interactome profiles were predicted for planarian Smed-CBP-2 and Smed-CBP-3 (Fig. 9 and S8). The interaction of both Smed-CBP-2 and Smed-CBP-3 with several proteins known to be regulated and/or directly acetylated by human CBP/p300 homologues appeared conserved in planarians (Fig. 9). Interestingly, some interactors such as p53, Runx1, or Ets2 have been already functionally characterized in planarians (see references listed in Fig. 8A) and their essential roles in processes such as neoblast proliferation, specification and differentiation might be related to their regulation by CBPs. Altogether, these results indicate that albeit being expressed in different cells, Smed-CBP-2 and Smed-CBP-3 share their network of protein interactors, suggesting that their diversified function might be instructed by the cellular environment in which *Smed-cbp-2* or *Smed-cbp-3* are embraced.

**Fig. 9.**
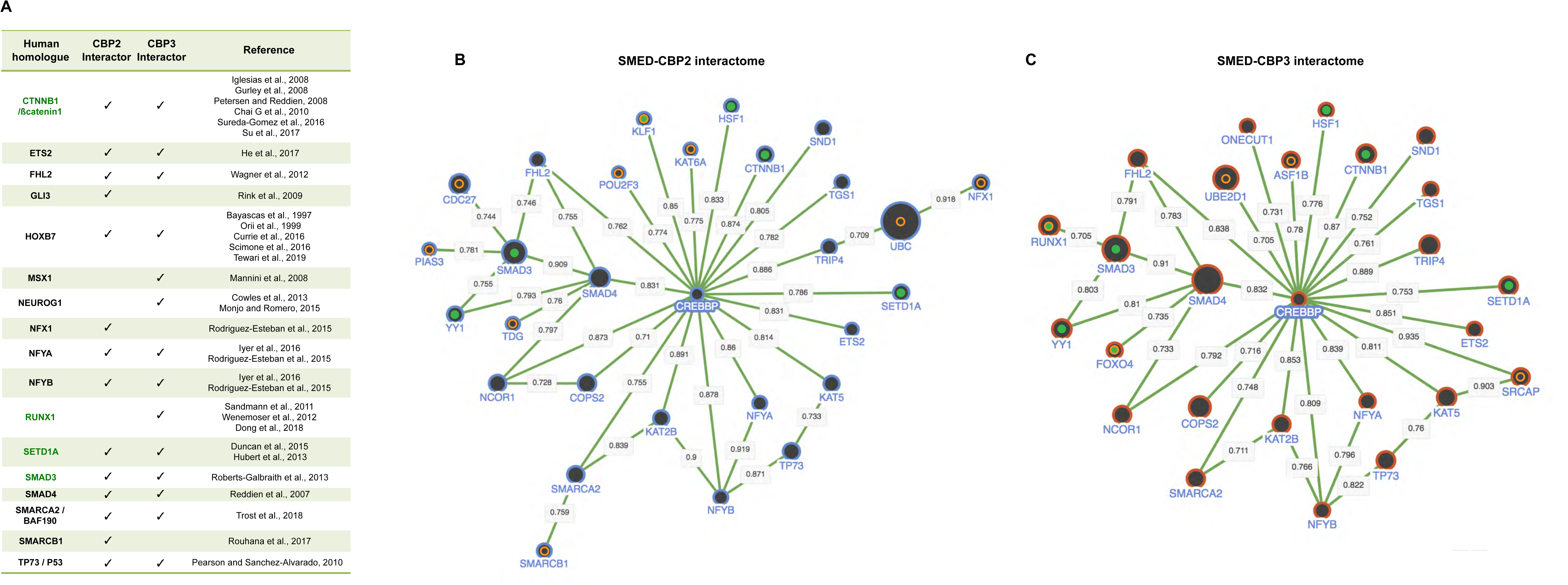
Smed-CBP-2 and Smed-CBP-3 predicted interactomes. (A) Summary table listing part of the interactors of Smed-CBP-2 and Smed-CBP-3 predicted by PlanNET (Castillo-Lara and Abril 2018). Only interactors that have been characterized in planarians are listed and the reference to the original data is recorded. Smed-CBPs interactors whose human homologs are acetylated by CREBBP are highlighted in green (Dancy and Cole 2015, https://thebiogrid.org, https://www.phosphosite.org/homeAction.action). Refer to Fig. S8 for the complete list of Smed-CBP-2 and Smed-CBP-3 predicted interactors. (B-C) Representation of the network of predicted interactions for Smed-CBP-2 (isotig 23520, counterpart of dd_Smed_v6_7719) (B) and Smed-CBP-3 (dd_Smed_v6_9343) (C) using the NetExplorer tool available at PlanNET website (Castillo-Lara and Abril 2018). Each node in the network represents a protein, the size of the nodes depends on the node degree (total number of interactions). Interactions are noted by lines between the proteins and numbers on the edges correspond to the measure of the confidence of any given interaction. Only protein connections with a confidence higher than 0,7 are represented. Interactors specific for each planarian homologue are highlighted with an orange circle. Proteins known to be acetylated by human CREBBP/p300 are highlighted with a green dot (Dancy and Cole 2015, https://thebiogrid.org, https://www.phosphosite.org/homeAction.action). For simplicity, the name of the human homologue protein is shown. Refer to tables in Fig. S8 for the corresponding numbers of the planarian homologue contigs.

## DISCUSSION

CREB-binding protein (CBP) and p300 have a central role in regulating gene expression in metazoans. CBP was originally characterized as a binding partner of CREB (cAMP-response element binding) protein (Chrivia et al., 1993). Their functions are mainly mediated by their role as transcriptional co-activators serving as scaffolds to bring together different factors at the promoter regions as well as through their lysine acetyltransferase activity. Thus, hundreds of interacting proteins have been described and dozens of proteins (histone and non-histone) have been shown to be acetylated by CBP/p300 proteins (Dancy and Cole, 2015; Holmqvist and Mannervik, 2013; Voss and Thomas, 2018). Whereas a single homologue of the CBP/p300 family has been identified in cnidarians, flies, molluscs and non-vertebrate chordates, a gene duplication occurred at the origin of vertebrates seems to be the origin of the CBP and p300 paralogs identified in vertebrates (Thomas and Kahn, 2016). Remarkably, this gene family has been expanded in Platyhelminthes. Thus, 2 homologues are found in the parasitic worm *Schistosoma mansoni* (Bertin et al., 2006) and 5 homologues have been identified in the freshwater planarians *Dendrocoelum lacteum* and *Schmidtea mediterranea* (Fig. S1). Other gene families such as those of *noggin* and *beta-catenin* are also expanded in planarians (Iglesias et al., 2008; Molina et al., 2009; Su et al., 2017).

Although vertebrates CBP and p300 show an extremely high degree of identity, they seem to have non-redundant functions (Thomas and Kahn, 2016). Thus, in the context of mammalian stem cell biology, for instance, CBP and p300 have been proposed to regulate proliferation and differentiation through their interaction with beta-catenin. A current model based on several results from a variety of progenitor/stem cells proposes that the interaction of beta-catenin with p300 would promote the expression of genes involved in cell differentiation whereas the interaction of beta-catenin with CBP would be required for stem cell renewal and maintenance (Thomas and Kahn, 2016; Manegold et al., 2018). Here, we have described 5 CBP homologues in *S. mediterranea* that seem to have originated from internal duplications in the Platyhelminthes lineage (Fig. S1). Planarian CBPs display distinct expression patterns based on whole-mount in situ hybridizations and single-cell expression (Fig. 1). Analyses at the single-cell level along the differentiation pathway for different cell lineages allow to distinguish between *Smed-cbp-1*, *-2* and *-3* and *Smed-cbp-4* and *-5*, as the expression of the first group increases, in general, from the neoblasts to fully differentiated cells, whereas the opposite is seen for *Smed-cbp-4* and *-5* (Fig. S2). The silencing of planarian cbp genes by RNAi have allowed us to characterize the function of *Smed-cbp-2* and *Smed-cbp-3*. Interestingly, these two genes seem to have opposite functions during planarian regeneration. Thus, the silencing of *Smed-cbp-2* perturbs regeneration and the planarians cannot grow normal blastemas. In these animals, there is a significant reduction in the neoblast population, *Smedwi-1* expression and the number of proliferating cells (Fig. 2). Despite these defects, neoblasts appear to be able to differentiate as even in the absence of normal blastemas photoreceptor cells and small brains differentiate around the wound region. Overall, many of the phenotypes observed after the silencing of *Smed-cbp-2* could be explained by a role of this gene in the maintenance and proliferation of the neoblasts. On the other side, the silencing of *Smed-cbp-3* results mainly in differentiation problems. After the silencing of *Smed-cbp-3* the animals grow normal blastemas but that remain mainly undifferentiated, with no external evidence of eye and body pigment cell differentiation. Internally, those blastemas contain an abnormally high number of neoblasts and the expression of neoblast markers such as *Smedwi-1* and *Smed-soxp-2* is also significantly increased (Fig. 6). Moreover, the differentiation of distinct neuronal populations, excretory cells and gut cells appear also highly impaired (Fig. 7). Based on these results it is tempting to speculate that *Smed-cbp-2* and *Smed-cbp-3* might have diverged functionally to regulate either stem cell maintenance and proliferation or differentiation in a similar way as it has been described for vertebrate CBP and p300. Whether or not these apparently opposite functions for planarians *cbp-2* and *cbp-3* could be mediated through the interaction of these factors with the Wnt/B-catenin pathway as it happens in mammalian stem cells remains for future analyses.

Although the silencing of *Smed-cbp-2* and *cbp-3* yields rather different phenotypes the expression patterns of these genes by in situ hybridizations (Fig. 1) and along the differentiation pathway for several cell lineages (Fig. S2) appears very similar. However, deep in silico analyses from an available planarian cell atlas (Plass et al., 2018) clearly indicates that the level of co-expression of these two genes is extremely low and quite similar in all cell lineages analyzed. These analyses also reveal that *Smed-cbp-3* is expressed at a higher level than *Smed-cbp-2* in all of the cell lineages and that both genes are expressed in progenitors and fully differentiated cells in all lineages, except for *Smed-cbp-2* in the protonephridia lineage (although here we only identified 4 cells expressing this gene) (Fig. S7). Also, for both genes their expression in the progenitor compartment (*Smedwi-1* positive) is slightly higher compared to the differentiated compartment (*Smedwi-1* negative) (Fig. S7). These results agree with the variety of phenotypes observed after the silencing of these genes in terms of the diversity of cell lineages affected and their probable requirement for progenitor specification as well as for the final differentiation of several cell types. As discussed above, vertebrate CBP/p300 proteins interact with hundreds of partners. However, the interactomes of CBP and p300 are in general very similar (Bedford et al., 2010; https://thebiogrid.org; https://string-db.org) suggesting that the different functions that these two genes may have are not probably caused by specific interactions with distinct sets of other proteins, but rather by the cellular context where these interactions may occur. Similarly, planarian *cbp-2* and *cbp-3* show very similar predicted interactomes (Fig. 9) that together with the extremely low level of co-expression further supports the idea that their functions may depend more on the cellular and molecular context than in their specific interacting partners.

Overall our results suggest that, in general, *cbp-2* might be involved in stem cell maintenance and proliferation whereas *cbp-3* might be required for the proper differentiation of several lineages. However, a more detailed analysis reveals that the silencing of each of these two genes does not affect in the same way the different cell lineages analyzed. Thus, whereas after the silencing of *cbp-2* there is a significant decrease in the *Smedwi-1+* population of stem cells that probably explains the subsequent decrease observed for epidermal, gut and photoreceptor progenitors, the neural progenitor compartment tends to increase (Fig. 5). This opens the possibility that *cbp-2* could have also a role in regulating the cell fate of planarian progenitors. This role would be similar to what it has been described in Xenopus embryos, where the inhibition of CBP/p300 function impairs the differentiation of non-neural tissues at the same time that induces neurogenesis throughout the entire embryo (Kato et al., 1999). Remarkably, the silencing of *cbp-2* results in the death of all the treated animals, both intact and regenerating, suggesting an additional role of *cbp-2* in cell survival. Previous studies have demonstrated the role of CBP on cell viability by inducing apoptosis after the inhibition of β-catenin/CBP signalling (Kleszcz et al., 2019). In the parasitic worm *Schistosoma mansoni* the silencing of the Smed-*cbp-2* homologue (called *Sm-cbp1,* Fig. S1) results also in a lethal phenotype apparently triggered by an increase in cell death (Collins and Collins, 2016). In *S. mansoni,* cell death triggered by Sm-cbp1 inhibition induces increased levels of neoblast proliferation that normally differentiate but that cannot survive because of the lack of function of *Sm-cbp1* (Collins and Collins, 2016). Similarly, although no increase in cell proliferation is observed after the silencing of *cbp-2*, there is a significant increment in cell death, suggesting that *Smed-cbp2* and *Sm-cbp1* could have a similar conserved role in cell survival.

On the other hand, the silencing of *cbp-3* affects cell differentiation in different ways depending on the cell lineage. Thus, after its inhibition, and despite the increased number of *Smedwi-1+* cells observed within the blastema, there is a high reduction in the number of neural, photoreceptor and protonephridia progenitors, but a concomitant increase in the number of epidermal and gut progenitors (Fig. 8), suggesting that *cbp-3* would regulate cell differentiation at different points along the pathway. Thus, *cbp-3* seems to be required for the specification of neural, photoreceptor and protonephridia progenitors but not for the specification of gut and epidermal progenitors. In these cases, *cbp-3* seems to be required for their final differentiation. The fact that different progenitor populations respond in opposite ways to the silencing of *cbp-3* further suggests that planarian *cbp* genes might have a role in cell fate determination.

As *cbp* genes may exert their function through multiple types of interactions with hundreds of factors it is difficult to determine how the silencing of *cbp-2* and *cbp-3* results in the high variety of phenotypes described here. Interestingly, however, the analyses of the predicted interactomes of these two proteins have identified putative interactions of CBP-2 and CBP-3 with multiple proteins that have been previously characterized in planarians (Fig. 9). Based on our current vertebrate knowledge some of these interactions would be mediated by the acetylation of partners such as B-CATENIN, RUNX1, SETD1A and SMAD3, whereas for partners such as ETS2, MSX1, NEUREGULIN and P53 other type of interactions would be established. The functional characterization of these factors allow us to suggest that some of their proposed functions might be mediated by CBP proteins. Thus, for example, the silencing of an ETS transcription factor results in a decrease in body-wall pigmentation (He et al., 2017). As described above the silencing of *cbp-3* leads to a failure to differentiate pigmented cells within the blastemas. Also, the silencing of *b-catenin* and *Runx1* has been linked to impaired neural differentiation (Sandmann et al., 2011; Sureda-Gómez et al., 2016; Dong et al., 2018; Zou et al., 2020). Similarly, CBP/p300 interact with Smad3 and p53 to regulate cell differentiation (Jain et al., 2012; Furumatsu et al., 2005; Martire et al. 2020). Finally, CBP/p300 have been shown to be required for normal epidermal regeneration through, for example, their interaction with KLF3 (Jones et al., 2020).

In summary, our results show that planarian *cbp* genes appear to have a conserved role as key regulators for the maintenance and differentiation of their stem cells (Fig. 10). As post-translational modifications play key roles in the regulation of stem cells proliferation and differentiation in humans and planarians (Strand et al., 2019; Wang et al., 2014) further experiments should determine which exact functions of *cbp-2* and *cbp-3* could be mediated either by the direct acetylation of their partners or by the acetylation of histone residues leading to chromatin remodeling also basic to regulate gene expression.

**Fig. 10.**
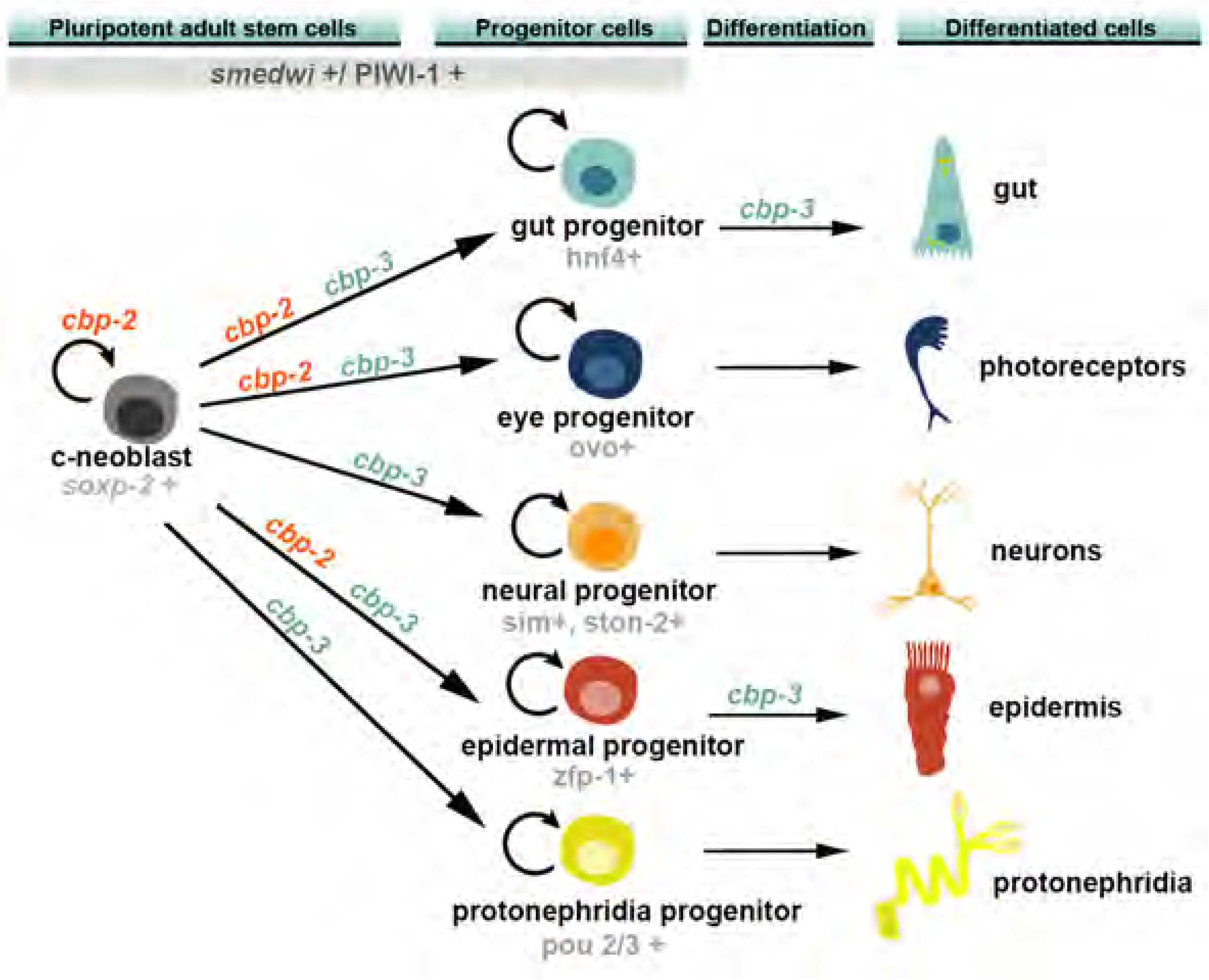
Proposed model of the function of *cbp-2* and *cbp-3* along the differentiation pathway of several cell lineages. See details within the text

## MATERIAL AND METHODS

### Animals and gene nomenclature

*Schmidtea mediterranea* from the asexual clonal line BCN-10 were used for all experiments. Animals were maintained at 20°C in PAM water 1x according to Cebrià and Newmark (2005). We fed the animals with veal ecological liver twice per week. Planarians were starved for at least one week before experiments.

### Sequence and protein domains arrangement analyses

*cbp* genes were identified from the *S. mediterranea* genome (Grohme et al., 2018) and amplified using specific primers (Table S1). Protein Domain conservation of *Smed-cbps* (Fig. 1C) was analyzed using SMART (http://smart.embl-heidelberg.de) and Pfam protein domain databases (http://pfam.xfam.org/).

### Phylogenetic analyses

Protein sequences of CBP/P300 homologues were obtained from NCBI and Planmine v3.0 (Rozanski et al., 2019) and aligned using MUSCLE. The evolutionary history was inferred using the Neighbor-Joining method conducted in MEGA X (Kumar et al., 2018; Stecher et al., 2020). Bootstrap values were determined by 1000 replicates and the rate variation among sites was modeled with a gamma distribution (shape parameter = 1). The evolutionary distances were computed using the Poisson correction method. The analysis involved 28 amino acid sequences. All ambiguous positions were removed for each sequence pair (pairwise deletion option) leading a total of 3817 positions in the final dataset. The tree was visualized using FigTree v1.4.4 (http://tree.bio.ed.ac.uk/software/figtree/).

### RNA Interference

Double-stranded RNA (dsRNA) for *Smed-cbp-1, -2, -3, -* and *-5* were synthesized as previously described (Sánchez-Alvarado and Newmark, 1999). Animals were injected during two rounds of 3 consecutive days each, with 4 days elapsed in between. Injections were done using the Nanoject II (Drummond Scientific, Broomall, PA, USA) and consisted of three injections of 32 nl of 1 µg/µl dsRNA per day. Control animals were injected with *gfp* dsRNA. One day after the second round of injection, planarians were amputated pre-pharyngeally to induce anterior regeneration. In some experiments, we kept the animals intact in order to analyze the effects of silencing the gene in homeostasis.

### Single-cell Sequencing (SCS) Data

*cbp* genes expression profiles were obtained via the planaria single-cell database hosted by Rajewsky lab at the Berlin Institute for Medical Systems Biology of the Max Delbrück Center in Berlin (Plass et al., 2018) and the Gene Co-expression Counts tool hosted in PlanNET website (Castillo-Lara and Abril 2018; Castillo-Lara et al., 2020).

### WISH and WFISH

Whole-mount *in situ* hybridization (WISH) was performed as previously described (Currie et al., 2016). Whole-mount fluorescent in situ hybridization (WFISH) was performed as previously described (King and Newmark, 2013). Riboprobes for in situ hybridization were synthesized using the DIG RNA labelling kit (Sp6/T7, Roche) following the manufacturer’s instructions. Samples were mounted in 70% glycerol/PBS solution.

### Immunohistochemistry

Whole-mount immunohistochemistry was performed as previously described (Ross et al., 2015). Treated animals were killed in cold 2% HCl in ultrapure H_2_O during 5 minutes and then washed with PBS-Tx (PBS + 0,3% Triton X-100) at RT shaking for 5 minutes. Then, they were put in a fixative solution (4% formaldehyde in PBS-Tx) for 15 minutes at RT with shaking and washed twice with PBS-Tx. Afterwards, samples were bleached in 6% H_2_O_2_ (in PBS-Tx) at RT during 16 hours under direct light. Next day, bleached animals were washed with PBS-Tx and incubated for 2 hours in 1% blocking solution (1% BSA in PBS-Tx), followed by the primary antibody (diluted in blocking solution) overnight at 4°C. We used as primary antibodies: anti-phospho-histone3 (PH3, Cell signalling technology) to detect mitotic cells which are between the G2 phase and the M phase (diluted 1/300); anti-SYNAPSIN used as a pan-neural marker (anti-SYNORF1 diluted 1/50, Developmental Studies Hybridoma Bank) ; anti-VC-1, specific for planarian photosensitive cells (1:15000; Sakai et al., 2000); anti-SMEDWI-1, specific for neoblasts diluted 1:1500; Guo et al., 2006; Marz et al., 2013); and TMUS-13, specific for myosin heavy chain (diluted 1/5; Cebrià et al., 1997); AA4.3, against a-tubulin to visualize the epithelial cilia (diluted 1/20, Developmental Studies Hybridoma Bank). Secondary antibodies used were: Alexa-488-conjugated goat anti-mouse diluted 1:400 (Molecular Probes) for SYNAPSIN, VC1, TMUS-13 and AA4.3 and Alexa 568-conjugated goat anti-rabbit diluted 1:1000 (Molecular Probes) for PH3 and SMEDWI-1. Samples were mounted in 70% glycerol/PBS solution. Nuclei were stained with DAPI (1:5000; Sigma-Aldrich) and TO-PRO®-3 (1:3000, Thermo Fisher Scientific, Waltham, MA, USA).

### Caspase-3 Activity assay

Caspase-3 activity was performed as described previously (González-Estévez et al., 2007) using 20 mg of protein extract, which was incubated for 2 h in the dark at 37°C with 20 µM caspase-3 substrate Ac-DEVD-AMC (BD Biosciences Pharmingen) or 2 ml from a stock of 1 mg/ml for a final volume of 150 µl. For each experiment, 3 biological replicates were made for both *gfp(RNAi)* and *Smed-cbp(RNAi)* conditions. For each condition, 5 planarians were used to do the protein extraction. Using a FLUOstar Optima Microplate Readers (BMG Labtech), the enzyme activity was measured in a luminescence spectrophotometer (Perkin-Elmer LS-50) (excitation: 380 nm; emission: 440 nm), describing one unit of caspase-3 activity as the amount of active enzyme necessary to produce an increase of one arbitrary luminescence unit after 2 hours of incubation.

### Microscopy, image acquisition and image analysis

Live animals were photographed with an sCM EX-3 High End Digital Microscope Camera DC.3000s (Visual Inspection Technology). WISH, WFISH and immunostained animals were observed with a stereomicroscope Leica MZ16F. Images were captured with the ProGres C3 camera from Jenoptik and then treated with Photoshop CS6 to mount the figures. Representative images of WFISH and immunostained animals were captured with confocal laser scanning microscopy (Leica TCS-SPE microscope) and treated with ImageJ1.51d and Photoshop CS6 to mount the figures.

### Quantitative real time PCR (qPCR)

qPCR was performed with three technical and biological replicates. Following RNAi injections, total RNA was isolated from a pool of 5 treated planarians per condition by homogenization in TRIzol Reagent (Invitrogen). Housekeeping gene ura4 was used to normalize the expression levels.

### Statistical analysis

All the comparisons are done with t-student test. Previously it was confirmed normality and homogeneity of the data with Shapiro-Wilk test and Bartlett test, respectively.

## SUPPORTING INFORMATION

**Fig. S1. CBP/P300 phylogenetic tree.** Evolutionary relationship of Smed-CBPs and CBP/p300 proteins reconstructed using the neighbor-joining method. The values beside the branches represent the percentage of times that a node was supported in 1000 bootstrap pseudoreplications. Only values higher than 70 are indicated. The scale indicates expected amino acid substitutions per site. The tree was rooted with the CBP homologue from *Arabidopsis*. The tree was rooted with the CBP homologue from Arabidopsis. Accession number of CBP/P300 protein sequences retrieved from NCBI are: *Arabidopsis thaliana* CBP AG60059.1; *Crassostrea gigas* CBP XP_011427370.2; *Drosophila melanogaster* CBP NP_001188576.1; *Gallus gallus* CBP XP_015150110.2 and P300 XP_004937767.1; *Homo sapiens* CBP NP_001073315.1 and P300 AAA18639.1; Hydra vulgaris CBP XP_002156492.2; *Mus musculus* CBP P45481.3 and P300 NP_808489.4; Nematostella vectensis CBP XP_032237806.1; *Octopus bimaculoides* CBP KOF77471.1; *Saccoglossus kowalevskii* CBP ADB22397.1; *Schistosoma mansoni* CBP1 AAZ57334.1 and CBP2 ABB17266.1; *Strongylocentrotus purpuratus* CBP XP_030836851.1; *Tribolium castaneum* CBP XP_008192363.1. Accession number of sequences retrieved from Planmine are: *Dendrocoelum lacteum* CBP-1 dd_Dlac_v9_182568_0_1; CBP-2 dd_Dlac_v9_190131_0_2; CBP-3 dd_Dlac_v9_194372_0_1; CBP-4 dd_Dlac_v9_191089_0_1; CBP-5 dd_Dlac_v9_185242_0_1; *Macrostomum lignano* CBP gr_Mlig_v2_1509_1063_1 and *Schmidtea mediterranea* CBP-1 dd_Smed_v6_8613_0_1; CBP-2 dd_Smed_v6_7719_0_1; CBP-3 dd_Smed_v6_9343_0_1; CBP-4 dd_Smed_v6_6340_0_1; CBP-5 dd_Smed_v6_16296_0_1.

**Fig. S2. Expression of *cbp* genes during differentiation of planarian main cell types.** Relative gene expression of *Smed-cbp* genes for the individual stages during differentiation of all major planarian cell types (in pseudotime) according to single cell analysis (https://shiny.mdc-berlin.de/psca/, Plass et al., 2018). *Smed-cbp-1*, *Smed-cbp-2* and *Smed-cbp-3* versus *Smed-cbp-4* and *Smed-cbp-5* expression have been grouped to facilitate comparison. a.u, arbitrary units.

**Fig. S3. *cbp-1*, -*4* and *-5* silencing does not impair either neoblast maintenance or differentiation during planarian regeneration.** Mitotic rate analysed by anti-PH3 antibody is not distorted after silencing *cbp-1*, -*4* and -*5*. At the same time, these treated animals do not show any apparent neoblast population impairment, after checking the *Smedwi-1* probe. Neural differentiation, analysed by both anti-SYNAPSIN antibody and *Smed-th* probe, seem to be not affected after silencing *cbp-1*, -*4* and -*5*. All animals were analysed at 10 days of regeneration. Scale bar: 200 µm.

**Fig. S4. Polarity markers are not affected after *cbp-2* silencing (A)** The posterior polarity marker *Smed-wntp-1* is not affected in regenerating heads after silencing *cbp-2.* **(B)** The anterior polarity marker *Smed-notum* is not affected after silencing *cbp-2* in regenerating trunks. All samples correspond to 7 days of regeneration. Scale bar: 200 µm.

**Fig. S5. *cbp-3* RNAi effects in intact planarians. (A)** Digestive and neural systems labelled by WISH for *Smed-pk* and immunostained with anti-SYNAPSIN, respectively, were not affected after silencing *cbp-3* in intact planarians. Also, neoblasts presented normal numbers and distribution in these animals. Mature epidermal cells labelled by *Smed-ifb* are absent from the most anterior region of intact planarians after silencing *cbp-3*. Scale bar: 200 µm. **(B)** Quantification of mitotic cells (PH3+) in intact planarians after silencing *cbp-3*. Values represent the mean ± s.e.m of an average of 5 samples per each condition. (*P< 0,05, Student’s t-test). **(C)** Combined neural progenitor *Smed-sim* WFISH and SMEDWI-1 immunostaining in homeostasis conditions showing the complete absence of these neural progenitors around the pharyngeal region. All images show intact animals after 29 days of RNAi treatment. Scale bar: 100 µm.

**Fig. S6. Differentiation defects in regenerating heads after silencing *cbp-3* (A)** In vivo images corresponding to regenerating heads after two rounds of injection and 10 days of regeneration. Note the lack of pigmentation in the anterior blastema. **(B)** WISH for *Smed-pk* digestive system and *Smed-laminin* pharynx markers in regenerating heads reveal impaired differentiation after *cbp-3* knockdown. **(C)** Several neural markers reveal severe defects in the regenerated brains after *cbp-3* silencing. All images correspond to regenerating heads after 10 days of regeneration. Scale bar: 200 µm.

**Fig. S7. *Smed-cbp-2* and *Smed-cbp-3* are expressed both in progenitor and differentiated cells of several planarian lineages but are barely co-expressed.** (**A**) Graph summarizing the % of cells of the main planarian cellular types expressing or co-expressing *Smed-cbp-2* and *Smed-cbp-3* genes. Note that less than 1,3% of the cells in each lineage co-express *Smed-cbp-2* and *Smed-cbp-3*. (B-F) *Left panels:* Relative gene expression of *Smed-cbp-2, Smed-cbp-3* and *Smedwi-1* for the individual stages during differentiation of neuronal (**B**), epidermal (**C**), muscular (**D**), intestinal (**E**) and excretory (**F**) cell types (in pseudotime) according to single cell analysis (Plass et al., 2018, https://shiny.mdc-berlin.de/psca/). a.u, arbitrary units. *Right panels:* Table presenting the number and percentage of cells expressing or co-expressing *Smed-cbp-2* and *Smed-cbp-3* genes in progenitors (*Piwi1*-positive) and differentiated (*Piwi1*-negative) cells of the neuronal (**B**), epidermal (**C**), muscular (**D**), intestinal (**E**) and excretory (**F**) lineages. **(A-F)** In all the analyses the number of cells expressing a given gene in each cellular type was obtained from the Gene Co-expression Counts tool of PlanEXP found in PlanNET (Castillo-Lara and Abril 2018; Castillo-Lara et al., 2020) using single cell data from Plass et al., 2018. As an example, in **A**, the percentage of cells expressing *Smed-cbp-2* in neoblasts was calculated by comparing the number of *Smed-cbp-2* expressing cells in this cellular type (434 cells) relative to the total number of cells categorized as neoblasts (8075 cells) in the single cell dataset. Similarly, the percentage of neural progenitor cells (**B**) expressing *Smed-cbp-2* was obtained by comparing the number of cells co-expressing *Smed-cbp-2* and *Smedwi-1* in the neural lineage related to the total number of *Smedwi-1*-positive cells categorized as neuronal in the single cell dataset.

**Fig. S8. Smed-CBP-2 and Smed-CBP-3 predicted interactomes** Tables listing the full Smed-CBP-2 (isotig 23520, counterpart of dd_Smed_v6_7719) and Smed-CBP-3 (dd_Smed_v6_9343) interactomes predicted by PlanNET (Castillo-Lara and Abril 2018). Only interactors with a confidence higher than 0,6 are listed. Interactors specific for each planarian CBP homologue are underlined and proteins that are known to be acetylated by human CREBBP/p300 are highlighted in green (Dancy and Cole 2015, https://thebiogrid.org, https://www.phosphosite.org). The reference to the original publication has been added for the homologues already characterized in planarians.

## Acknowledgments

We thank all members of the E. Saló and T. Adell laboratory for discussions; H. Orii and Prof. K. Watanabe for providing anti-VC-1; Monoclonal anti-SYNORF1 antibody was obtained from the Developmental Studies Hybridoma Bank, developed under the auspices of the National Institute of Child Health and Human Development and maintained by the Department of Biological Sciences, University of Iowa, Iowa City, IA, USA. MDM has received funding from the postdoctoral fellowship programme Beatriu de Pinós, funded by the Secretary of Universities and Research (Government of Catalonia) and by the Horizon 2020 programme of research and innovation of the European Union under the Marie Sklodowska-Curie grant agreement No 801370. FC was supported by grants BFU2015-65704-P and PGC2018-100747-B-100 from Ministerio de Ciencia, Innovación y Universidades, Spain and grant 2017 SGR 1455 from AGAUR, Generalitat de Catalunya.

## Author contributions

Conceived and designed the experiments: SF, MDM, FC. Performed the experiments: SF, SC, CV, JG, JM. Analyzed the data: SF, SC, CV, JG, MDM, JM, FC, RR, TS, KB. Wrote the paper: SF, MDM, FC.

## References

Avgustinova, A. and S. A. Benitah (2016). “Epigenetic control of adult stem cell function.” Nat Rev Mol Cell Biol 17(10): 643–658. doi: 10.1038/nrm.2016.76

Baguñà, J., E. Saló, J. Collet, M. C. Auladell & M. Ribas, 1988. Cellular, molecular and genetic approaches to regeneration and pattern formation in planarians. Fortschr. Zool. 36: 65–78.

Baguñà, J. (2012) The planarian neoblast: the rambling history of its origin and some current black boxes. Int J Dev Biol 56(1-3):19–37. doi: 10.1387/ijdb.113463jb.

Bayascas, J. R., Castillo, E., Muñoz-Mármol, A. M. and Saló, E. (1997). “Planarian Hox genes: novel patterns of expression during regeneration.” Development 124(1): 141–148.

Bedford, D. C., Kasper, L. H., Fukuyama, T. and Brindle, P. K. (2010). “Target gene context influences the transcriptional requirement for the KAT3 family of CBP and p300 histone acetyltransferases.” Epigenetics 5(1): 9–15. doi: 10.4161/epi.5.1.10449

Bertin B, Oger F, Cornette J, Caby S, Noël C, Capron M, Fantappie MR, Rumjanek FD and Pierce RJ. (2006) Schistosoma mansoni CBP/p300 has a conserved domain structure and interacts functionally with the nuclear receptor SmFtz-F1. Mol Biochem Parasitol. 146(2):180–91. doi: 10.1016/j.molbiopara.2005.12.006.

Bonuccelli, L., Rossi, L., Lena, A., Scarcelli, V., Rainaldi, G., Evangelista, M., Iacopetti, P., Gremigni, V. and Salvetti, A. (2010). “An RbAp48-like gene regulates adult stem cells in planarians.” J Cell Sci 123(Pt 5): 690–698. doi: 10.1242/jcs.053900

Brai, E., Marathe, S., Astori, S., Fredj, N. B., Perry, E., Lamy, C., Scotti, A. and Alberi, L. (2015). “Notch1 Regulates Hippocampal Plasticity Through Interaction with the Reelin Pathway, Glutamatergic Transmission and CREB Signaling.” Front Cell Neurosci 9: 447. doi: 10.3389/fncel.2015.00447

Castillo-Lara, S. and J. F. Abril (2018). “PlanNET: homology-based predicted interactome for multiple planarian transcriptomes.” Bioinformatics 34(6): 1016–1023. doi: 10.1093/bioinformatics/btx738

Castillo-Lara, S., Pascual-Carreras, E. and Abril, J. F. (2020). “PlanExp: intuitive integration of complex RNA-seq datasets with planarian omics resources.” Bioinformatics 36(6): 1889–1895. doi: 10.1093/bioinformatics/btz802

Cebrià, F., Vispo, M., Newmark, P., Bueno, D. and Romero, R. (1997). “Myocyte differentiation and body wall muscle regeneration in the planarian Girardia tigrina.” Dev Genes Evol 207(5): 306–316. doi: 10.1007/s004270050118

Cebrià, F. and Newmark, P.A. (2005). Planarian homologs of netrin and netrin receptor are required for proper regeneration of the central nervous system and the maintenance of nervous system architecture. Development 132(16):3691–703. doi: 10.1242/dev.01941.

Chai, G., Ma, C., Bao, K., Zheng, L., Wang, X., Sun, Z., Salò, E., Adell, T. and Wu, W. (2010). “Complete functional segregation of planarian beta-catenin-1 and -2 in mediating Wnt signaling and cell adhesion.” J Biol Chem 285(31): 24120–24130. doi: 10.1074/jbc.M110.113662

Chan, H. M. and N. B. La Thangue (2001). “p300/CBP proteins: HATs for transcriptional bridges and scaffolds.” J Cell Sci 114(Pt 13): 2363–2373.

Chandebois, R. (1980). “Cell sociology and the problem of automation in the development of pluricellular animals.” Acta Biotheor 29(1): 1–35. doi: 10.1007/BF00045880

Cheng, L. C., Tu, K. C., Seidel, C. W., Robb, S. M. C., Guo, F. and Sánchez Alvarado, A. (2018). “Cellular, ultrastructural and molecular analyses of epidermal cell development in the planarian Schmidtea mediterranea.” Dev Biol 433(2): 357–373. doi: 10.1016/j.ydbio.2017.08.030

Chrivia, J. C.,Kwok, R. P., Lamb, N., Hagiwara, M., Montminy, M. R. and Goodman, R. H. (1993). “Phosphorylated CREB binds specifically to the nuclear protein CBP.” Nature 365(6449): 855–859. doi: 10.1038/365855a0

Collins, J. N. and J. J. Collins, 3rd (2016). “Tissue Degeneration following Loss of Schistosoma mansoni cbp1 Is Associated with Increased Stem Cell Proliferation and Parasite Death In Vivo.” PLoS Pathog 12(11): e1005963. doi: 10.1371/journal.ppat.1005963

Cowles, M. W., Brown, D. D., Nisperos, S. V., Stanley, B. N., Pearson, B. J. and Zayas, R. M. (2013). “Genome-wide analysis of the bHLH gene family in planarians identifies factors required for adult neurogenesis and neuronal regeneration.” Development 140(23): 4691–4702. doi: 10.1242/dev.098616

Currie, K. W., Brown, D. D., Zhu, S., Xu, C., Voisin, V., Bader, G. D. and Pearson, B. J. (2016). “HOX gene complement and expression in the planarian Schmidtea mediterranea.” Evodevo 7: 7. doi: 10.1186/s13227-016-0044-8

Dancy, B. M. and P. A. Cole (2015). “Protein lysine acetylation by p300/CBP.” Chem Rev 115(6): 2419–2452. doi: 10.1021/cr500452k

Dattani A, Kao D, Mihaylova Y, Abnave P, Hughes S, Lai A, Sahu S and Aboobaker AA. Epigenetic analyses of planarian stem cells demonstrate conservation of bivalent histone modifications in animal stem cells. Genome Res. 28(10):1543–1554. doi: 10.1101/gr.239848.118.

Dattani, A., Sridhar, D. and Aziz Aboobaker, A. (2019). “Planarian flatworms as a new model system for understanding the epigenetic regulation of stem cell pluripotency and differentiation.” Semin Cell Dev Biol 87: 79–94. doi: 10.1016/j.semcdb.2018.04.007

Dong, Z., Yang, Y., Chen, G. and Liu, D. (2018). “Identification of runt family genes involved in planarian regeneration and tissue homeostasis.” Gene Expr Patterns 29: 24–31. doi: 10.1016/j.gep.2018.04.006

Duncan, E. M., Chitsazan, A. D., Seidel, C. W. and Alvarado, A. S. (2016). “Set1 and MLL1/2 Target Distinct Sets of Functionally Different Genomic Loci In Vivo.” Cell Rep 17(3): 930. doi: 10.1016/j.celrep.2016.09.071

Dutto, I., Scalera, C. and Prosperi, E. (2018). “CREBBP and p300 lysine acetyl transferases in the DNA damage response.” Cell Mol Life Sci 75(8): 1325–1338. doi: 10.1007/s00018-017-2717-4

Esvald, E. E., Tuvikene, J., Sirp, A., Patil, S., Bramham, C. R. and Timmusk, T. (2020). “CREB Family Transcription Factors Are Major Mediators of BDNF Transcriptional Autoregulation in Cortical Neurons.” J Neurosci 40(7): 1405–1426. doi: 10.1523/jneurosci.0367-19.2019

Fincher, C. T., Wurtzel, O., de Hoog, T., Kravarik, K. M. and Reddien, P. W. (2018). “Cell type transcriptome atlas for the planarian Schmidtea mediterranea.” Science 360(6391). Doi: 10.1126/science.aaq1736

Furumatsu, T., Tsuda, M., Taniguchi, N., Tajima, Y. and Asahara, H. (2005). “Smad3 induces chondrogenesis through the activation of SOX9 via CREB-binding protein/p300 recruitment.” J Biol Chem 280(9): 8343–8350. Doi: 10.1074/jbc.M413913200

Gavino, M. A. and P. W. Reddien (2011). “A Bmp/Admp regulatory circuit controls maintenance and regeneration of dorsal-ventral polarity in planarians.” Curr Biol 21(4): 294–299. doi: 10.1016/j.cub.2011.01.017

Giordano, A. and M. L. Avantaggiati (1999). “p300 and CBP: partners for life and death.” J Cell Physiol 181(2): 218–230. doi: 10.1002/(sici)1097-4652(199911)181:2<218::Aid-jcp4>3.0.Co;2-5

Godini, R., Lafta, H. Y. and Fallahi, H. (2018). “Epigenetic modifications in the embryonic and induced pluripotent stem cells.” Gene Expr Patterns 29: 1–9. doi: 10.1016/j.gep.2018.04.001

Goodman, R. H. and S. Smolik (2000). “CBP/p300 in cell growth, transformation, and development.” Genes Dev 14(13): 1553–1577.

González-Estévez, C., Felix, D. A., Aboobaker, A. A. and Saló, E. (2007). “Gtdap-1 promotes autophagy and is required for planarian remodeling during regeneration and starvation.” Proc Natl Acad Sci U S A 104(33): 13373–13378. doi: 10.1073/pnas.0703588104

Grohme, M. A., Schloissnig, S., Rozanski, A., Pippel, M., Young, G. R., Winkler, S., Brandl, H., Henry, I., Dahl, A., Powell, S., Hiller, M., Myers, E. and Rink, J. C. (2018). “The genome of Schmidtea mediterranea and the evolution of core cellular mechanisms.” Nature 554(7690): 56–61. doi: 10.1038/nature25473

Guo L, Zhang S, Rubinstein B, Ross E and Sánchez Alvarado, A. (2006). Widespread maintenance of genome heterozygosity in Schmidtea mediterranea. Nat Ecol Evol 1(1):19. doi: 10.1038/s41559-016-0019.

He, X., Lindsay-Mosher, N., Li, Y., Molinaro, A. M., Pellettieri, J. and Pearson, B. J. (2017). “FOX and ETS family transcription factors regulate the pigment cell lineage in planarians.” Development 144(24): 4540–4551. doi: 10.1242/dev.156349

Hardingham, G. E., Arnold, F. J. and Bading, H. (2001). “Nuclear calcium signaling controls CREB-mediated gene expression triggered by synaptic activity.” Nat Neurosci 4(3): 261–267. doi: 10.1038/85109

He, X., Lindsay-Mosher, N., Li,Y., Molinaro, A. M., Pellettieri, J., and Pearson B. J. (2017) FOX and ETS family transcription factors regulate the pigment cell lineage in planarians. Development 144(24): 4540–4551. doi: 10.1242/dev.156349

Henderson, J. M., Nisperos, S. V., Weeks, J., Ghulam, M., Marín, I. and Zayas, R. M. (2015). “Identification of HECT E3 ubiquitin ligase family genes involved in stem cell regulation and regeneration in planarians.” Dev Biol 404(2): 21–34. doi: 10.1016/j.ydbio.2015.04.021

Holmqvist, P. H. and M. Mannervik (2013). “Genomic occupancy of the transcriptional co-activators p300 and CBP.” Transcription 4(1): 18–23. doi: 10.4161/trns.22601

Hubert, A., Henderson, J. M., Ross, K. G., Cowles, M. W., Torres, J. and Zayas, R. M. (2013). “Epigenetic regulation of planarian stem cells by the SET1/MLL family of histone methyltransferases.” Epigenetics 8(1): 79–91. doi: 10.4161/epi.23211

Hyman, L. H. (1951) The invertebrates : Platyhelminthes and Rhynchocoela, the acoelomate Bilateria. New York : McGraw-Hill Book Company Inc. Vol.2 pp.vii + 550 pp.

Iglesias, M., Gomez-Skarmeta, J. L., Salo, E., Adell, T. (2008). “Silencing of Smed-betacatenin1 generates radial-like hypercephalized planarians.” Development 135(7): 1215–1221. doi: 10.1242/dev.020289

Iyer, H., Collins, J. J., 3rd and Newmark, P. A. (2016). “NF-YB Regulates Spermatogonial Stem Cell Self-Renewal and Proliferation in the Planarian Schmidtea mediterranea.” PLoS Genet 12(6): e1006109. doi: 10.1371/journal.pgen.1006109

Jain, A. K., Allton, K., Iacovino, M., Mahen, E., Milczarek, R. J., Zwaka, T. P., Kyba, M. and Barton, M. C. (2012). “p53 regulates cell cycle and microRNAs to promote differentiation of human embryonic stem cells.” PLoS Biol 10(2): e1001268. Doi: 10.1371/journal.pbio.1001268

Jones, J., Chen, Y., Tiwari, M., Li, J., Ling, J. and Sen, G. L. (2020). “KLF3 Mediates Epidermal Differentiation through the Epigenomic Writer CBP.” Science 23(7): 101320. doi: 10.1016/j.isci.2020.101320

Kato, Y., Shi, Y. and He, X. (1999). “Neuralization of the Xenopus embryo by inhibition of p300/ CREB-binding protein function.” J Neurosci 19(21): 9364–9373. doi: 10.1523/jneurosci.19-21-09364.1999

King, R. S. and P. A. Newmark (2013). “In situ hybridization protocol for enhanced detection of gene expression in the planarian Schmidtea mediterranea.” BMC Dev Biol 13: 8. doi: 10.1186/1471-213x-13-8

Kleszcz, R., Szymańska, A., Krajka-Kuźniak, V., Baer-Dubowska, W. and Paluszczak, J. (2019). “Inhibition of CBP/β-catenin and porcupine attenuates Wnt signaling and induces apoptosis in head and neck carcinoma cells.” Cell Oncol (Dordr) 42(4): 505–520. doi: 10.1007/s13402-019-00440-4

Kumar S., Stecher G., Li M., Knyaz C., and Tamura K. (2018). MEGA X: Molecular Evolutionary Genetics Analysis across computing platforms. Molecular Biology and Evolution 35:1547–1549.

Lapan, S. W. and P. W. Reddien (2012). “Transcriptome analysis of the planarian eye identifies ovo as a specific regulator of eye regeneration.” Cell Rep 2(2): 294–307. doi: 10.1016/j.celrep.2012.06.018

Lundblad, J. R., Kwok, R. P., Laurance, M. E., Harter, M. L. and Goodman, R. H. (1995). “Adenoviral E1A-associated protein p300 as a functional homologue of the transcriptional co-activator CBP.” Nature 374(6517): 85–88. doi: 10.1038/374085a0

Manegold, P., Lai, K. K. Y., Wu, Y., Teo, J. L., Lenz, H. J., Genyk, Y. S., Pandol, S. J., Wu, K., Lin, D. P., Chen, Y., Nguyen, C., Zhao, Y. and Kahn, M. (2018). “Differentiation Therapy Targeting the β-Catenin/CBP Interaction in Pancreatic Cancer.” Cancers (Basel) 10(4). Doi: 10.3390/cancers10040095

Mannini, L., Deri, P., Gremigni, V., Rossi, L., Salvetti, A. and Batistoni, R. (2008). “Two msh/msx-related genes, Djmsh1 and Djmsh2, contribute to the early blastema growth during planarian head regeneration.” Int J Dev Biol 52(7): 943–952. doi: 10.1387/ijdb.072476lm

Martire, S., Nguyen, J., Sundaresan, A. and Banaszynski, L. A. (2020). “Differential contribution of p300 and CBP to regulatory element acetylation in mESCs.” BMC Mol Cell Biol 21(1): 55. doi: 10.1186/s12860-020-00296-9

März, M., Seebeck, F. and Bartscherer, K. (2013). A Pitx transcription factor controls the establishment and maintenance of the serotonergic lineage in planarians. Development. 140(22):4499–509. doi: 10.1242/dev.100081.

Mayr, B. and M. Montminy (2001). “Transcriptional regulation by the phosphorylation-dependent factor CREB.” Nat Rev Mol Cell Biol 2(8): 599–609. doi: 10.1038/35085068

Mihaylova, Y., Abnave, P., Kao, D., Hughes, S., Lai, A., Jaber-Hijazi, F., Kosaka, N. and Aboobaker, A. A. (2018). “Conservation of epigenetic regulation by the MLL3/4 tumour suppressor in planarian pluripotent stem cells.” Nat Commun 9(1): 3633. doi: 10.1038/s41467-018-06092-6

Molina, M. D., Salo, E. and Cebria, F. (2007). “The BMP pathway is essential for re-specification and maintenance of the dorsoventral axis in regenerating and intact planarians.” Dev Biol 311(1): 79–94. doi:10.1016/j.ydbio.2007.08.019

Molina, M. D., Saló, E. and Cebrià, F. (2009). “Expression pattern of the expanded noggin gene family in the planarian Schmidtea mediterranea.” Gene Expr Patterns 9(4): 246–253. doi: 10.1016/j.gep.2008.12.008

Molinaro, A. M. and B. J. Pearson (2016). “In silico lineage tracing through single cell transcriptomics identifies a neural stem cell population in planarians.” Genome Biol 17: 87. doi: 10.1186/s13059-016-0937-9

Orii, H., Kato, K., Umesono, Y., Sakurai, T., Agata, K. and Watanabe, K. (1999). “The planarian HOM/HOX homeobox genes (Plox) expressed along the anteroposterior axis.” Dev Biol 210(2): 456–468. doi: 10.1006/dbio.1999.9275

Owlarn, S., Klenner, F., Schmidt, D., Rabert, F., Tomasso, A., Reuter, H., Mulaw, M. A., Moritz, S., Gentile, L., Weidinger, G. and Bartscherer, K. (2017). “Generic wound signals initiate regeneration in missing-tissue contexts.” Nat Commun 8(1): 2282. doi: 10.1038/s41467-017-02338-x

Pearson, B. J. and Sánchez Alvarado, A. (2010). “A planarian p53 homolog regulates proliferation and self-renewal in adult stem cell lineages.” Development 137(2): 213–221. doi: 10.1242/dev.044297

Pellettieri, J., Fitzgerald, P., Watanabe, S., Mancuso, J., Green, D.R. and Sánchez Alvarado, A (2010). “Cell death and tissue remodeling in planarian regeneration.” Dev Biol 338(1): 76–85. ddoi: 10.1016/j.ydbio.2009.09.015

Petersen, C. P. and P. W. Reddien (2008). “Smed-betacatenin-1 is required for anteroposterior blastema polarity in planarian regeneration.” Science 319(5861): 327–330. doi: 10.1126/science.1149943

Plass, M., Solana, J., Wolf, F. A., Ayoub, S., Misios, A., Glažar, P., Obermayer, B., Theis, F. J., Kocks, C. and Rajewsky, N. (2018). “Cell type atlas and lineage tree of a whole complex animal by single-cell transcriptomics.” Science 360(6391). doi: 10.1126/science.aaq1723

Reddien, P. W., Bermange, A. L., Kicza, A. M. and Sánchez Alvarado, A. (2007). “BMP signaling regulates the dorsal planarian midline and is needed for asymmetric regeneration.” Development 134(22): 4043–4051. doi: 10.1242/dev.007138

Reddien, P. W. (2013). “Specialized progenitors and regeneration.” Development 140(5): 951–957. doi: 10.1242/dev.080499

Reddien, P. W. (2018). “The Cellular and Molecular Basis for Planarian Regeneration.” Cell 175(2): 327–345. doi: 10.1016/j.cell.2018.09.021

Rink, J. C., Gurley, K. A., Elliott, S. A. and Sánchez Alvarado, A. (2009). “Planarian Hh signaling regulates regeneration polarity and links Hh pathway evolution to cilia.” Science 326(5958): 1406–1410. doi: 10.1126/science.1178712

Rink, J. C. (2013). “Stem cell systems and regeneration in planaria.” Dev Genes Evol 223(1-2): 67–84. doi: 10.1007/s00427-012-0426-4

Roberts-Galbraith, R. H. and P. A. Newmark (2013). “Follistatin antagonizes activin signaling and acts with notum to direct planarian head regeneration.” Proc Natl Acad Sci U S A 110(4): 1363–1368. doi: 10.1073/pnas.1214053110

Robb, S. M. and Sánchez Alvarado, A. (2014). “Histone modifications and regeneration in the planarian Schmidtea mediterranea.” Curr Top Dev Biol 108: 71–93. doi: 10.1016/b978-0-12-391498-9.00004-8

Rodríguez-Esteban, G., González-Sastre, A., Rojo-Laguna, J. I., Saló, E. and Abril, J. F. (2015). “Digital gene expression approach over multiple RNA-Seq data sets to detect neoblast transcriptional changes in Schmidtea mediterranea.” BMC Genomics 16(1): 361. doi: 10.1186/s12864-015-1533-1

Rompolas, P., Patel-King, R. S. and King, S. M. (2010). “An outer arm Dynein conformational switch is required for metachronal synchrony of motile cilia in planaria.” Mol Biol Cell 21(21): 3669–3679. doi: 10.1091/mbc.E10-04-0373

Ross, K. G., Omuro, K. C., Taylor, M. R., Munday, R. K., Hubert, A., King, R. S. and Zayas, R. M. (2015). “Novel monoclonal antibodies to study tissue regeneration in planarians.” BMC Dev Biol 15: 2. doi: 10.1186/s12861-014-0050-9

Rouhana, L., Tasaki, J., Saberi, A. and Newmark, P. A. (2017). “Genetic dissection of the planarian reproductive system through characterization of Schmidtea mediterranea CPEB homologs.” Dev Biol 426(1): 43–55. doi: 10.1016/j.ydbio.2017.04.008

Rozanski, A., Moon, H., Brandl, H., Martín-Durán, J. M., Grohme, M. A., Hüttner, K., Bartscherer, K., Henry, I. and Rink, J. C. (2019). “PlanMine 3.0-improvements to a mineable resource of flatworm biology and biodiversity.” Nucleic Acids Res 47(D1): D812–d820. doi: 10.1093/nar/gky1070

Sakai, F., Agata, K., Orii, H. and Watanabe, K. (2000). Organization and regeneration ability of spontaneous supernumerary eyes in planarians -eye regeneration field and pathway selection by optic nerves-. Zoolog Sci 17(3):375–81. doi: 10.2108/jsz.17.375.

Salo, E. and J. Baguñá (1984). “Regeneration and pattern formation in planarians. I. The pattern of mitosis in anterior and posterior regeneration in Dugesia (G) tigrina, and a new proposal for blastema formation.” J Embryol Exp Morphol 83: 63–80.

Sánchez Alvarado, A. and P. A. Newmark (1999). “Double-stranded RNA specifically disrupts gene expression during planarian regeneration.” Proc Natl Acad Sci U S A 96(9): 5049–5054. doi: 10.1073/pnas.96.9.5049

Sandmann, T., Vogg, M. C., Owlarn, S., Boutros, M. and Bartscherer, K. (2011). “The head-regeneration transcriptome of the planarian Schmidtea mediterranea.” Genome Biol 12(8): R76. doi: 10.1186/gb-2011-12-8-r76

Scimone, M. L., Meisel, J. and Reddien, P. W. (2010). “The Mi-2-like Smed-CHD4 gene is required for stem cell differentiation in the planarian Schmidtea mediterranea.” Development 137(8): 1231–1241. doi: 10.1242/dev.042051

Scimone, M. L., Srivastava, M., Bell, G. W. and Reddien, P. W. (2011). “A regulatory program for excretory system regeneration in planarians.” Development 138(20): 4387–4398. doi: 10.1242/dev.068098

Scimone, M. L., Kravarik, K. M., Lapan, S. W. and Reddien, P. W. (2014). “Neoblast specialization in regeneration of the planarian Schmidtea mediterranea.” Stem Cell Reports 3(2): 339–352. doi: 10.1016/j.stemcr.2014.06.001

Scimone, M. L., Cote, L. E., Rogers, T. and Reddien, P. W. (2016). “Two FGFRL-Wnt circuits organize the planarian anteroposterior axis.” Elife 5. doi: 10.7554/eLife.12845

Shaywitz, A. J. and M. E. Greenberg (1999). “CREB: a stimulus-induced transcription factor activated by a diverse array of extracellular signals.” Annu Rev Biochem 68: 821–861. doi: 10.1146/annurev.biochem.68.1.821

Stecher G., Tamura K., and Kumar S. (2020). Molecular Evolutionary Genetics Analysis (MEGA) for macOS. Molecular Biology and Evolution. 37(4): 1237–1239. doi: 10.1093/molbev/msz312

Strand, N. S., Allen, J. M. and Zayas, R. M. (2019). “Post-translational regulation of planarian regeneration.” Semin Cell Dev Biol 87: 58–68. doi: 10.1016/j.semcdb.2018.04.009

Su, H., Sureda-Gomez, M., Rabaneda-Lombarte, N., Gelabert, M., Xie, J., Wu, W. and Adell, T. (2017). “A C-terminally truncated form of β-catenin acts as a novel regulator of Wnt/β-catenin signaling in planarians.” PLoS Genet 13(10): e1007030. doi: 10.1371/journal.pgen.1007030

Sureda-Gómez, M., Martín-Durán, J. M. and Adell, T. (2016). “Localization of planarian β-CATENIN-1 reveals multiple roles during anterior-posterior regeneration and organogenesis.” Development 143(22): 4149–4160. doi: 10.1242/dev.135152

Tewari, A. G., Owen, J. H., Petersen, C. P., Wagner, D. E. and Reddien, P. W. (2019). “A small set of conserved genes, including sp5 and Hox, are activated by Wnt signaling in the posterior of planarians and acoels.” PLoS Genet 15(10): e1008401. doi: 10.1371/journal.pgen.1008401

Thiruvalluvan, M., Barghouth, P. G., Tsur, A., Broday, L. and Oviedo, N. J. (2018). “SUMOylation controls stem cell proliferation and regional cell death through Hedgehog signaling in planarians.” Cell Mol Life Sci 75(7): 1285–1301. doi: 10.1007/s00018-017-2697-4

Thomas, P. D. and M. Kahn (2016). “Kat3 coactivators in somatic stem cells and cancer stem cells: biological roles, evolution, and pharmacologic manipulation.” Cell Biol Toxicol 32(1): 61–81. doi: 10.1007/s10565-016-9318-0

Trost, T., Haines, J., Dillon, A., Mersman, B., Robbins, M., Thomas, P. and Hubert, A. (2018). “Characterizing the role of SWI/SNF-related chromatin remodeling complexes in planarian regeneration and stem cell function.” Stem Cell Res 32: 91–103. doi: 10.1016/j.scr.2018.09.004

van Wolfswinkel, J. C., Wagner, D. E. and Reddien, P. W. (2014). “Single-cell analysis reveals functionally distinct classes within the planarian stem cell compartment.” Cell Stem Cell 15(3): 326–339. doi: 10.1016/j.stem.2014.06.007

Vásquez-Doorman, C. and C. P. Petersen (2016). “The NuRD complex component p66 suppresses photoreceptor neuron regeneration in planarians.” Regeneration (Oxf) 3(3): 168–178. doi: 10.1002/reg2.58

Voss, A. K. and T. Thomas (2018). “Histone Lysine and Genomic Targets of Histone Acetyltransferases in Mammals.” Bioessays 40(10): e1800078. doi: 10.1002/bies.201800078

Wagner, D. E., Wang, I. E. and Reddien, P. W. (2011). “Clonogenic neoblasts are pluripotent adult stem cells that underlie planarian regeneration.” Science 332(6031): 811–816. doi:10.1126/science.1203983

Wagner, D. E., Ho, J. J. and Reddien, P. W. (2012). “Genetic regulators of a pluripotent adult stem cell system in planarians identified by RNAi and clonal analysis.” Cell Stem Cell 10(3): 299–311. doi: 10.1016/j.stem.2012.01.016

Wang, F., Marshall, C. B. and Ikura, M. (2013). “Transcriptional/epigenetic regulator CBP/p300 in tumorigenesis: structural and functional versatility in target recognition.” Cell Mol Life Sci 70(21): 3989–4008. doi: 10.1007/s00018-012-1254-4

Wang, Y. C., Peterson, S. E. and Loring, J. F.(2014). “Protein post-translational modifications and regulation of pluripotency in human stem cells.” Cell Res 24(2): 143–160. doi: 10.1038/cr.2013.151

Wen, A. Y., Sakamoto, K. M. and Miller, L. S. (2010). “The role of the transcription factor CREB in immune function.” J Immunol 185(11): 6413–6419. doi: 10.4049/jimmunol.1001829

Wenemoser, D., Lapan, S. W.,Wilkinson, A. W., Bell, G. W. And Reddien, P. W. (2012). “A molecular wound response program associated with regeneration initiation in planarians.” Genes Dev 26(9): 988–1002. doi: 10.1101/gad.187377.112

Wenemoser, D. and P. W. Reddien (2010). “Planarian regeneration involves distinct stem cell responses to wounds and tissue absence.” Dev Biol 344(2): 979–991. doi: 10.1016/j.ydbio.2010.06.017

Wurtzel, O., Oderberg, I. M. and Reddien, P. W. (2017). “Planarian Epidermal Stem Cells Respond to Positional Cues to Promote Cell-Type Diversity.” Dev Cell 40(5): 491–504.e495. doi: 10.1016/j.devcel.2017.02.008

Yuan, L. W. and A. Giordano (2002). “Acetyltransferase machinery conserved in p300/CBP-family proteins.” Oncogene 21(14): 2253–2260. doi: 10.1038/sj.onc.1205283

Zeng A, Li H, Guo L, Gao X, McKinney S, Wang Y, Yu Z, Park J, Semerad C, Ross E, Cheng LC, Davies E, Lei K, Wang W, Perera A, Hall K, Peak A, Box A, Sánchez Alvarado A. (2018). Prospectively Isolated Tetraspanin(+) Neoblasts Are Adult Pluripotent Stem Cells Underlying Planaria Regeneration. Cell. 173(7):1593–1608.e20. doi: 10.1016/j.cell.2018.05.006.

Zou, L., Li, H., Han, X., Qin, J. and Song, G. (2020). “Runx1t1 promotes the neuronal differentiation in rat hippocampus.” Stem Cell Res Ther 11(1): 160. doi: 10.1186/s13287-020-01667-x

Zhu, S. J., Hallows, S. E., Currie, K. W., Xu, C. and Pearson, B. J. (2015). “A mex3 homolog is required for differentiation during planarian stem cell lineage development.” Elife 4. doi: 10.7554/eLife.07025

Zhu, L. and Pearson, B.J. (2016). (Neo)blast from the past: new insights into planarian stem cell lineages. Curr Opin Genet Dev 40:74–80. doi: 10.1016/j.gde.2016.06.007.

Zhu, S. J. and B. J. Pearson (2018). “Smed-myb-1 Specifies Early Temporal Identity during Planarian Epidermal Differentiation.” Cell Rep 25(1): 38–46.e33. doi: 10.1016/j.celrep.2018.09.011

Zuckerkandl, E. and L. Pauling (1965). “Molecules as documents of evolutionary history.” J Theor Biol 8(2): 357–366. doi: 10.1016/0022-5193(65)90083-4.

