## Supplemental Figures for "Planarian CREB-binding protein (CBP) gene family regulates stem cell maintenance and differentiation"

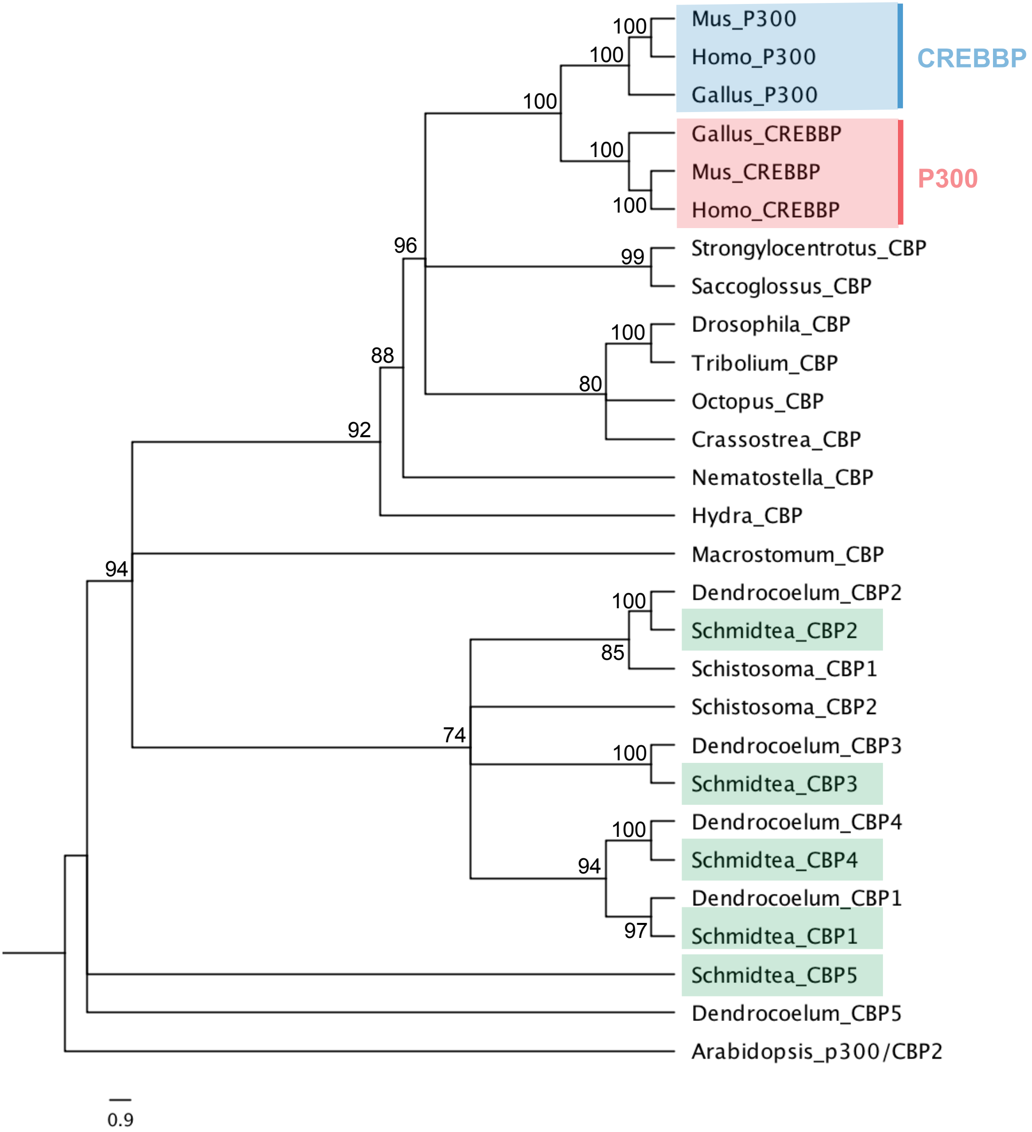
Mus_P300
Homo_P300
Gallus_P300
CREBBP
Gallus_CREBBP
Mus_CREBBP
Homo_CREBBP
P300
Strongylocentrotus_CBP
Saccoglossus_CBP
Drosophila_CBP
Tribolium_CBP
Octopus_CBP
Crassostrea_CBP
Nematostella_CBP
Hydra_CBP
Macrostomum_CBP
Dendrocoelum_CBP2
Schmidtea_CBP2
Schistosoma_CBP1
Schistosoma_CBP2
Dendrocoelum_CBP3
Schmidtea_CBP3
Dendrocoelum_CBP4
Schmidtea_CBP4
Dendrocoelum_CBP1
Schmidtea_CBP1
Schmidtea_CBP5
Dendrocoelum_CBP5
Arabidopsis_p300/CBP2
100
100
100
100
100
96
99
100
88
80
92
94
100
85
74
100
100
94
97
0.9

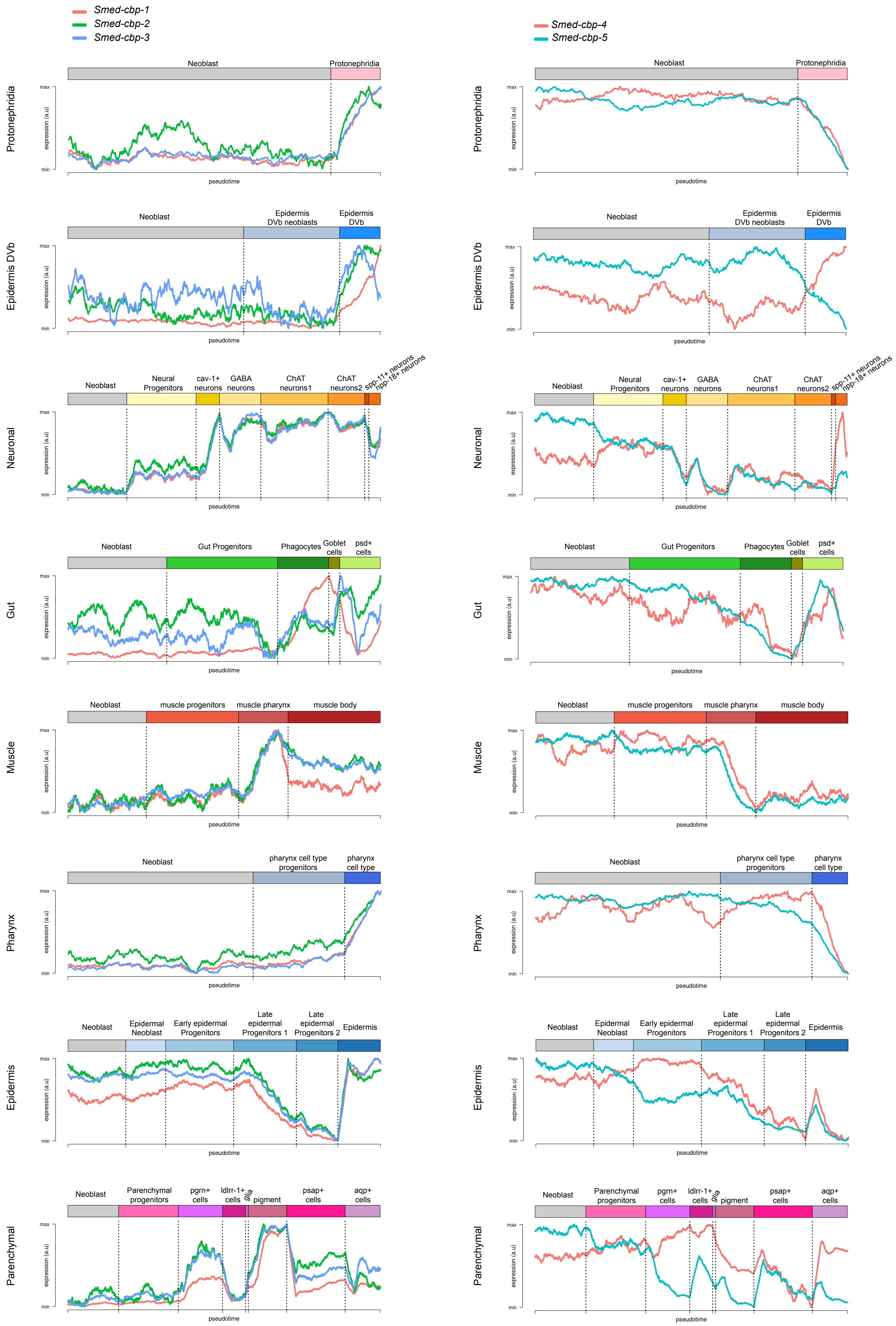
Smed-cbp-1
Smed-cbp-2
Smed-cbp-3
Smed-cbp-4
Smed-cbp-5
Protonephridia
Neoblast
Protonephridia
max
min
expression (a.u)
pseudotime
Epidermis DVb
Neoblast
Epidermis DVb neoblasts
Epidermis DVb
Neuronal
Neoblast
Neural Progenitors
cav-1+ neurons
GABA neurons
ChAT neurons1
ChAT neurons2
spp-11+ neurons
npp-18+ neurons
Gut
Neoblast
Gut Progenitors
Phagocytes
Goblet cells
psd+ cells
Muscle
Neoblast
muscle progenitors
muscle pharynx
muscle body
Pharynx
Neoblast
pharynx cell type progenitors
pharynx cell type
Epidermis
Neoblast
Epidermal Neoblast
Early epidermal Progenitors
Late epidermal Progenitors 1
Late epidermal Progenitors 2
Epidermis
Parenchymal
Neoblast
Parenchymal progenitors
pgrn+ cells
ldlrr-1+ cells
glia
pigment
psap+ cells
aqp+ cells

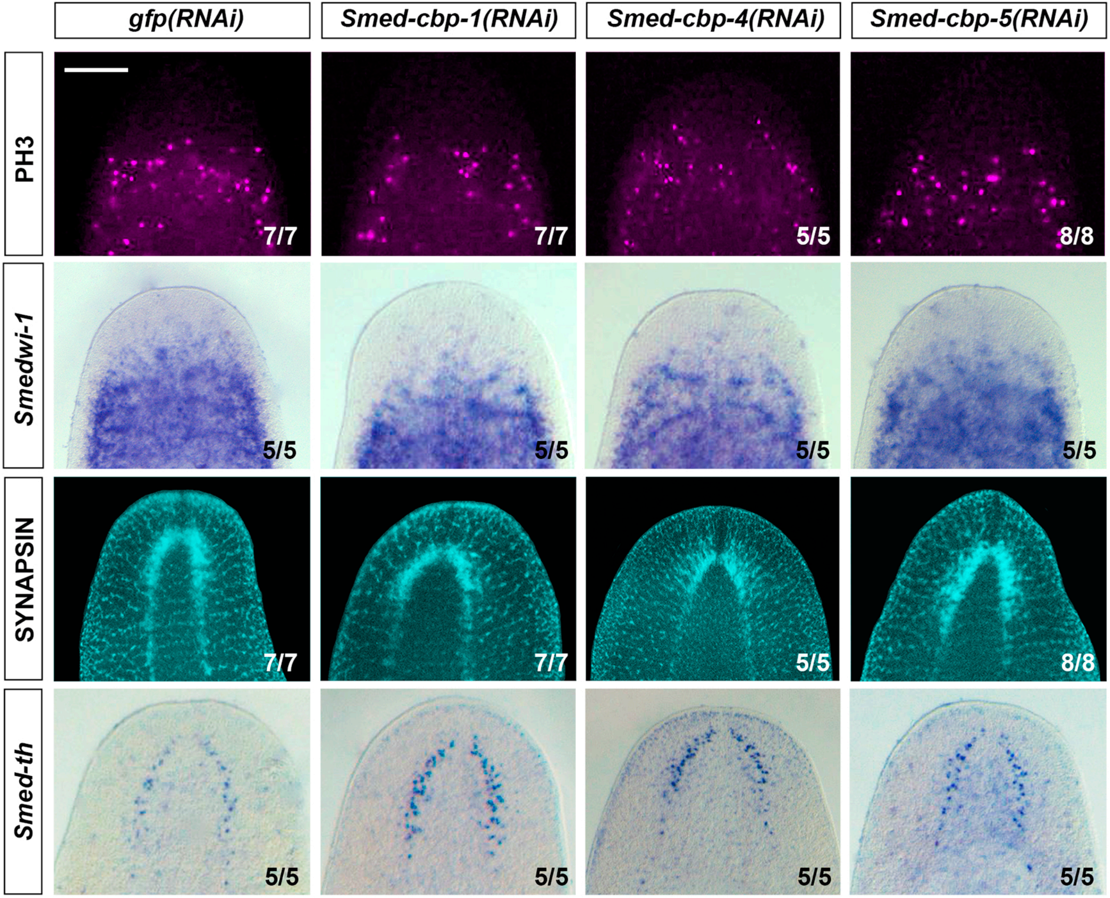

gfp(RNAi)
Smed-cbp-1(RNAi)
Smed-cbp-4(RNAi)
Smed-cbp-5(RNAi)
PH3
Smedwi-1
SYNAPSIN
Smed-th
7/7
7/7
5/5
8/8
5/5
5/5
5/5
5/5
7/7
7/7
5/5
8/8
5/5
5/5
5/5
5/5

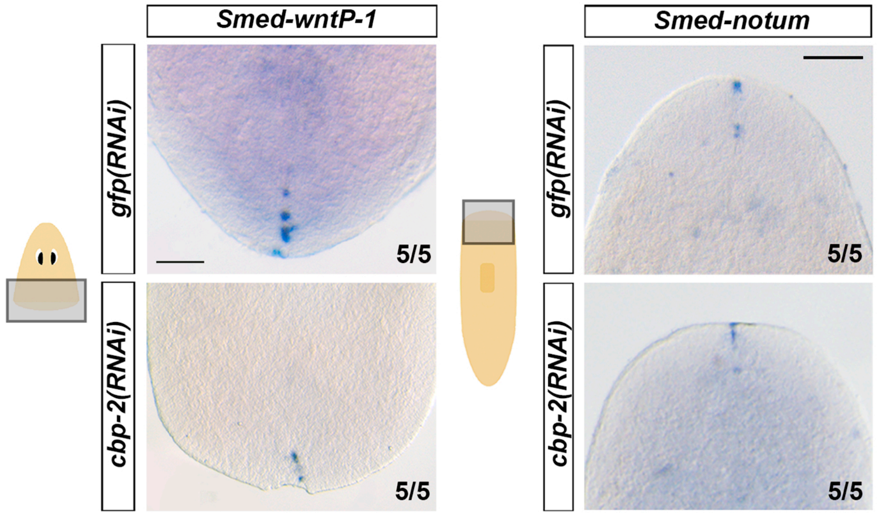
Smed-wntP-1
Smed-notum
gfp(RNAi)
cbp-2(RNAi)
5/5
5/5
5/5
5/5

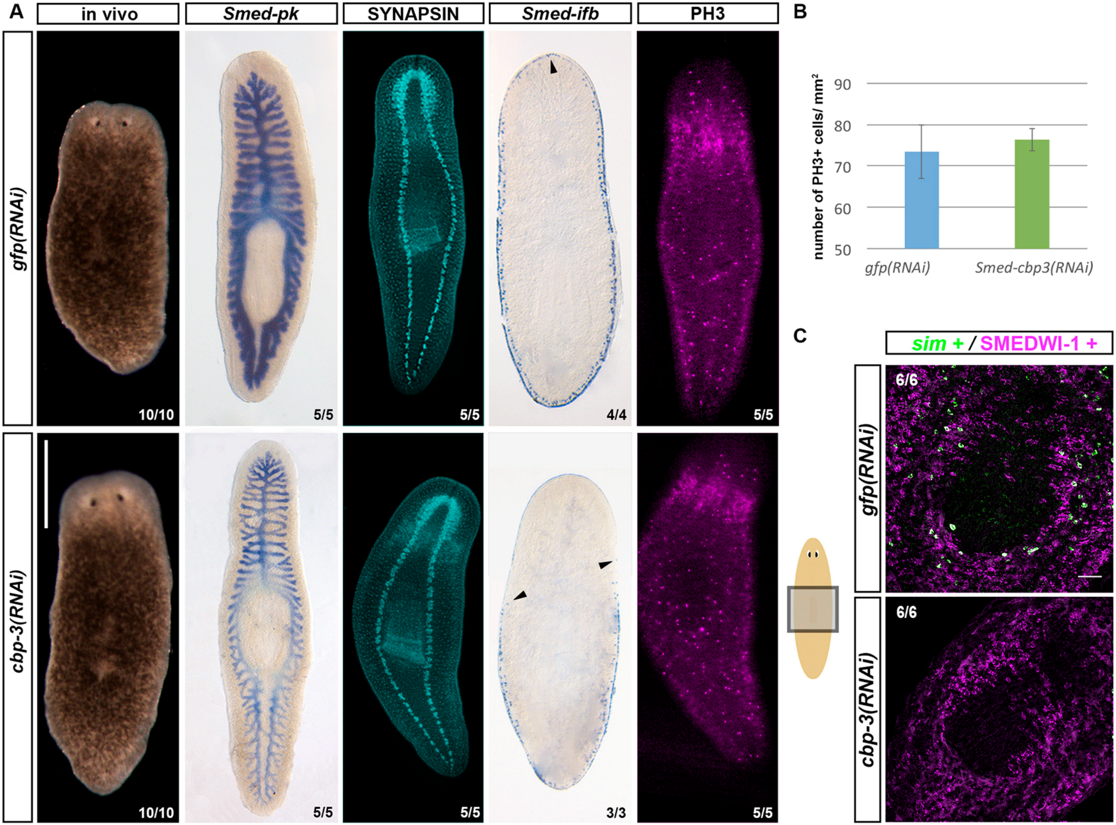

A
in vivo
Smed-pk
SYNAPSIN
Smed-ifb
PH3
gfp(RNAi)
cbp-3(RNAi)
10/10
5/5
5/5
4/4
5/5
10/10
5/5
5/5
3/3
5/5
B
number of PH3+ cells/ mm²
90
80
70
60
50
gfp(RNAi)
Smed-cbp3(RNAi)
C
sim + / SMEDWI-1 +
gfp(RNAi)
cbp-3(RNAi)
6/6
6/6

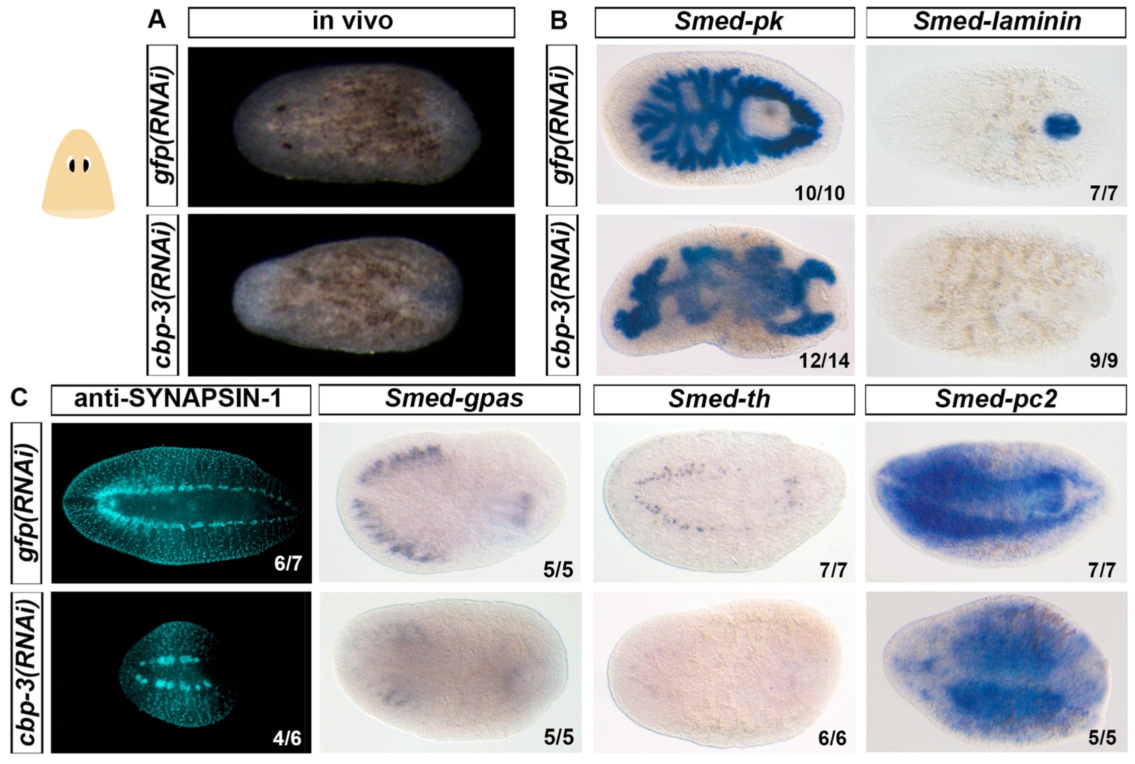
A
in vivo
gfp(RNAi)
cbp-3(RNAi)
B
Smed-pk
Smed-laminin
10/10
7/7
12/14
9/9
C
anti-SYNAPSIN-1
Smed-gpas
Smed-th
Smed-pc2
6/7
5/5
7/7
7/7
4/6
5/5
6/6
5/5

A

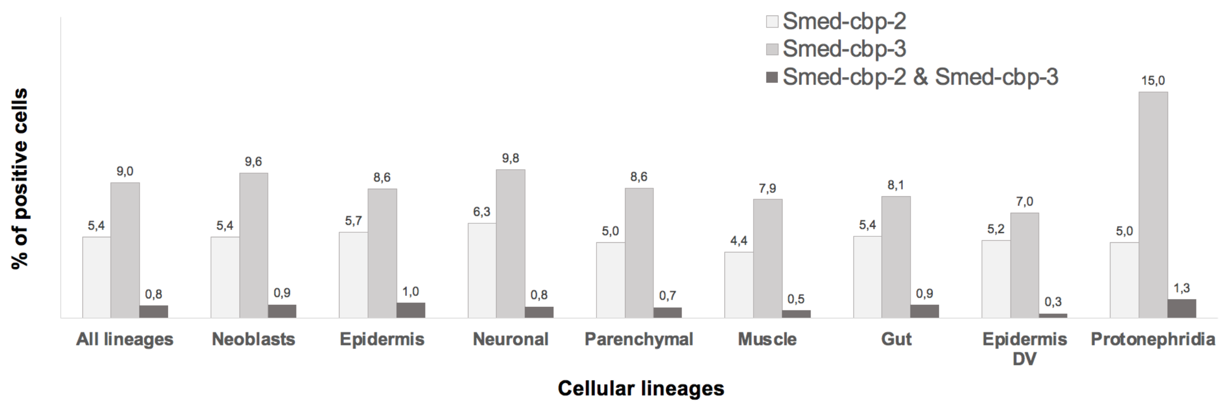


B

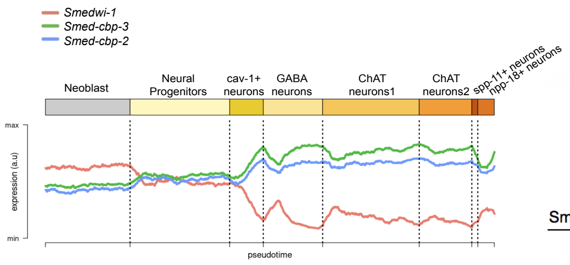


| | | Number of cells | | % of cells | |
|---|---|---|---|---|---|
| | | Progenitors (Piwi +) | Differentiated (Piwi -) | Progenitors (Piwi +) | Differentiated (Piwi -) |
| Smed-cbp-2 | 148 | 49 | 99 | 10,8 | 5,2 |
| Smed-cbp-3 | 232 | 58 | 174 | 12,8 | 9,1 |
| Smed-cbp-2 & Smed-cbp-3 | 18 | 7 | 11 | 1,5 | 0,6 |
| # cells neural lineage | 2358 | 453 | 1905 | | |

C

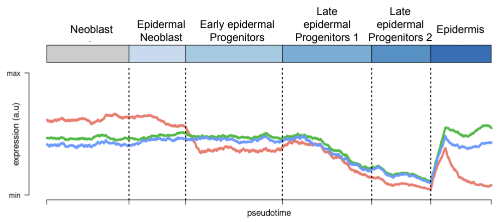


| | | Number of cells | | % of cells | |
|---|---|---|---|---|---|
| | | Progenitors (Piwi +) | Differentiated (Piwi -) | Progenitors (Piwi +) | Differentiated (Piwi -) |
| Smed-cbp-2 | 228 | 38 | 190 | 7,6 | 5,4 |
| Smed-cbp-3 | 344 | 52 | 292 | 10,4 | 8,3 |
| Smed-cbp-2 & Smed-cbp-3 | 40 | 7 | 33 | 1,4 | 0,9 |
| # cells epidermal lineage | 4009 | 498 | 3511 | | |

D

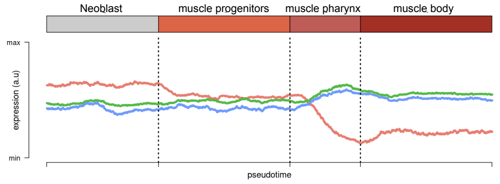


| | | Number of cells | | % of cells | |
|---|---|---|---|---|---|
| | | Progenitors (Piwi +) | Differentiated (Piwi -) | Progenitors (Piwi +) | Differentiated (Piwi -) |
| Smed-cbp-2 | 103 | 29 | 74 | 7,1 | 3,8 |
| Smed-cbp-3 | 185 | 60 | 125 | 14,7 | 6,4 |
| Smed-cbp-2 & Smed-cbp-3 | 12 | 7 | 5 | 1,7 | 0,3 |
| # cells muscular lineage | 2351 | 407 | 1944 | | |

E

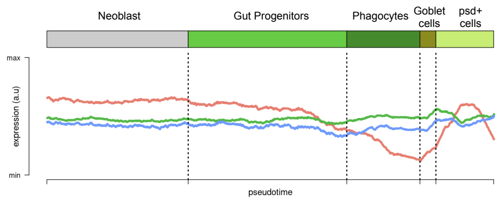


| | | Number of cells | | % of cells | |
|---|---|---|---|---|---|
| | | Progenitors (Piwi +) | Differentiated (Piwi -) | Progenitors (Piwi +) | Differentiated (Piwi -) |
| Smed-cbp-2 | 50 | 24 | 26 | 7,8 | 4,2 |
| Smed-cbp-3 | 74 | 29 | 45 | 9,5 | 7,3 |
| Smed-cbp-2 & Smed-cbp-3 | 8 | 1 | 7 | 0,3 | 1,1 |
| # cells gut lineage | 919 | 306 | 613 | | |

F

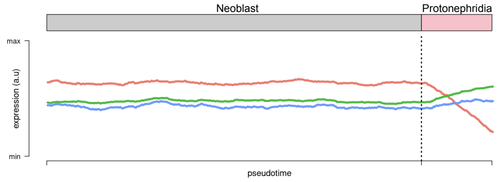


| | | Number of cells | | % of cells | |
|---|---|---|---|---|---|
| | | Progenitors (Piwi +) | Differentiated (Piwi -) | Progenitors (Piwi +) | Differentiated (Piwi -) |
| Smed-cbp-2 | 4 | 0 | 4 | 0,0 | 5,4 |
| Smed-cbp-3 | 12 | 2 | 10 | 33,3 | 13,5 |
| Smed-cbp-2 & Smed-cbp-3 | 1 | 0 | 1 | 0,0 | 1,4 |
| # cells protonephridia lineage | 80 | 6 | 74 | | |

### Smed-CBP-2 Interactome

| Human homologue | Planarian homologue | Confidence | Reference |
|---|---|---|---|
| CDC27 | isotig22501 | 0.643 | |
| COPS2 | isotig15449 | 0.71 | |
| CPSF4 | isotig01551 | 0.656 | |
| CTNNB1 /ßcatenin1 | isotig23001<br>isotig22585 | 0.874<br>0.83 | Iglesias et al., 2008<br>Gurley et al., 2008<br>Petersen and Reddien, 2008<br>Chai G et al., 2010<br>Sureda-Gomez et al., 2016<br>Su et al., 2017 |
| EP300 | isotig22472 | 0.609 | |
| ETS2 | isotig12474<br>isotig12473 | 0.831<br>0.798 | He et al., 2017 |
| FHL2 | isotig25875<br>isotig15616<br>isotig21493<br>isotig13394<br>isotig13395 | 0.796<br>0.762<br>0.762<br>0.79<br>0.766 | Wagner et al., 2012 |
| GLI3 | isotig22707 | 0.614 | Rink et al., 2009 |
| HIPK2 | isotig21321 | 0.626 | |
| HOXB7 | isotig18915 | 0.61 | Bayascas et al., 1997<br>Orii et al., 1999<br>Currie et al., 2016<br>Scimone et al., 2016<br>Tewari et al., 2019 |
| HSF1 | isotig25963<br>isotig09567 | 0.833<br>0.791 | |
| HTT | isotig14108 | 0.663 | |
| KAT2B | isotig22939 | 0.891 | |
| KAT5 | isotig09000 | 0.814 | |
| KAT6A | isotig23440 | 0.775 | |
| KLF1 | isotig20465 | 0.85 | |
| NCOR1 | isotig23154 | 0.873 | |
| NFX1 | isotig13021 | 0.613 | Rodriguez-Esteban et al., 2015 |
| NFYA | isotig24638<br>isotig24612 | 0.86<br>0.816 | Iyer et al., 2016<br>Rodriguez-Esteban et al., 2015 |
| NFYB | isotig18238<br>isotig21277 | 0.878<br>0.82 | Iyer et al., 2016<br>Rodriguez-Esteban et al., 2015 |
| ONECUT1 | isotig25733<br>isotig22450 | 0.645<br>0.608 | |
| PIAS3 | isotig22235<br>isotig01445 | 0.608<br>0.608 | |
| POU2F3 | isotig21518 | 0.774 | |
| PPP2R5E | isotig14006 | 0.623 | |
| SETD1A | isotig19076 | 0.786 | Duncan et al., 2015<br>Hubert et al., 2013 |
| SMAD3 | isotig13864 | 0.612 | Roberts-Galbraith et al., 2013 |
| SMAD4 | isotig25984 | 0.831 | Reddien et al., 2007 |
| SMARCA2 / BAF190 | isotig22778 | 0.755 | Trost et al., 2018 |
| SMARCB1 | isotig11019 | 0.608 | Rouhana et al., 2017 |
| SND1 | isotig13514 | 0.805 | |
| TDG | isotig23364 | 0.604 | |
| TGS1 | isotig07164<br>isotig11458<br>isotig11459 | 0.782<br>0.755<br>0.738 | |
| TP73 / P53 | isotig17795 | 0.643 | Pearson and Sanchez-Alvarado, 2010 |
| TRIP4 | isotig23604 | 0.886 | |
| UBC | isotig00999 | 0.66 | |
| YY1 | isotig14820 | 0.607 | |

### Smed-CBP-3 Interactome

| Human homologue | Planarian homologue | Confidence | Reference |
|---|---|---|---|
| ASF1B | dd_Smed_v6_5120_0_1 | 0.78 | |
| BRPF1 | dd_Smed_v6_9528_0_1 | 0.6 | |
| CDC27 | dd_Smed_v6_7556_0_1 | 0.687 | |
| COPS2 | dd_Smed_v6_4989_0_1 | 0.716 | |
| CPSF4 | dd_Smed_v6_4069_0_1<br>dd_Smed_v6_24321_0_1 | 0.678<br>0.657 | |
| CTBP1 | dd_Smed_v6_43551_0_1<br>dd_Smed_v6_25893_0_1 | 0.692<br>0.692 | |
| CTNNB1 /ßcatenin1 | dd_Smed_v6_2688_0_1<br>dd_Smed_v6_4850_0_1<br>dd_Smed_v6_9667_0_1 | 0.87<br>0.847<br>0.776 | Iglesias et al., 2008<br>Gurley et al., 2008<br>Petersen and Reddien, 2008<br>Chai G et al., 2010<br>Sureda-Gomez et al., 2016<br>Su et al., 2017 |
| DYRK1B | dd_Smed_v6_3773_0_1<br>dd_Smed_v6_6670_0_1 | 0.617<br>0.603 | |
| ETS2 | dd_Smed_v6_2092_0_1 | 0.851 | He et al., 2017 |
| FHL2 | dd_Smed_v6_5037_0_1<br>dd_Smed_v6_5014_0_1<br>dd_Smed_v6_5854_0_1<br>dd_Smed_v6_2771_0_1<br>dd_Smed_v6_34269_0_1 | 0.839<br>0.838<br>0.805<br>0.805<br>0.797 | Wagner et al., 2012 |
| FOXO4 | dd_Smed_v6_3040_0_1 | 0.608 | |
| HIPK2 | dd_Smed_v6_7724_0_1 | 0.607 | |
| HOXB7 | dd_Smed_v6_22524_0_1<br>dd_Smed_v6_16227_0_1 | 0.677<br>0.658 | Bayascas et al., 1997<br>Orii et al., 1999<br>Currie et al., 2016<br>Scimone et al., 2016<br>Tewari et al., 2019 |
| HSF1 | dd_Smed_v6_7535_0_1<br>dd_Smed_v6_9099_0_2 | 0.825<br>0.776 | |
| HTT | dd_Smed_v6_3049_0_1 | 0.668 | |
| KAT2B | dd_Smed_v6_11453_0_1<br>dd_Smed_v6_12274_0_1 | 0.853<br>0.786 | |
| KAT5 | dd_Smed_v6_4706_1_1 | 0.811 | |
| MSX1 | dd_Smed_v6_18505_0_1 | 0.603 | Mannini et al., 2008 |
| NCOR1 | dd_Smed_v6_7709_0_1 | 0.792 | |
| NEUROG1 | dd_Smed_v6_9906_0_1<br>dd_Smed_v6_26877_0_1 | 0.646<br>0.646 | Cowles et al., 2013<br>Monjo and Romero, 2015 |
| NFYA | dd_Smed_v6_4860_0_1<br>dd_Smed_v6_18122_0_1<br>dd_Smed_v6_8585_0_1 | 0.867<br>0.839<br>0.82 | Iyer et al., 2016<br>Rodriguez-Esteban et al., 2015 |
| NFYB | dd_Smed_v6_5828_0_1 | 0.809 | Iyer et al., 2016<br>Rodriguez-Esteban et al., 2015 |
| ONECUT1 | dd_Smed_v6_16472_0_1<br>dd_Smed_v6_25197_0_1<br>dd_Smed_v6_7877_0_1 | 0.731<br>0.672<br>0.672 | |
| PSMC5 | dd_Smed_v6_1176_0_1 | 0.608 | |
| RUNX1 | dd_Smed_v6_3565_0_2 | 0.651 | Sandmann et al., 2011<br>Wenemoser et al., 2012<br>Dong et al., 2018 |
| SETD1A | dd_Smed_v6_9988_0_1 | 0.753 | Duncan et al., 2015<br>Hubert et al., 2013 |
| SMAD2 | dd_Smed_v6_8193_0_1 | 0.673 | Roberts-Galbraith et al., 2013 |
| SMAD4 | dd_Smed_v6_1923_0_1<br>dd_Smed_v6_19757_0_1 | 0.832<br>0.774 | Reddien et al., 2007 |
| SMARCA2 / BAF190 | dd_Smed_v6_16980_0_1 | 0.748 | Trost et al., 2018 |
| SND1 | dd_Smed_v6_906_0_1 | 0.752 | |
| SRCAP | dd_Smed_v6_8618_0_1<br>dd_Smed_v6_12273_0_1<br>dd_Smed_v6_14019_0_1 | 0.935<br>0.904<br>0.886 | |
| TGS1 | dd_Smed_v6_11559_0_1<br>dd_Smed_v6_9949_0_1 | 0.761<br>0.734 | |
| TP73 / p53 | dd_Smed_v6_5563_0_1 | 0.698 | Pearson and Sanchez-Alvarado, 2010 |
| TRIP4 | dd_Smed_v6_9211_0_1 | 0.889 | |
| UBE2D1 | dd_Smed_v6_6773_0_1 | 0.705 | |
| YY1 | dd_Smed_v6_10092_0_1 | 0.61 | |
